# Shedding light on the underlying characteristics of genomes using Kronecker model families of codon evolution

**DOI:** 10.1101/2020.08.12.247890

**Authors:** Maryam Zaheri, Nicolas Salamin

## Abstract

The mechanistic models of codon evolution rely on some simplistic assumptions in order to reduce the computational complexity of estimating the high number of parameters of the models. This paper is an attempt to investigate how much these simplistic assumptions are misleading when they violate the nature of the biological dataset in hand. We particularly focus on three simplistic assumptions made by most of the current mechanistic codon models including: 1) only single substitutions between nucleotides within codons in the codon transition rate matrix are allowed. 2) mutation is homogenous across nucleotides within a codon. 3) assuming HKY nucleotide model is good enough at the nucleotide level. For this purpose, we developed a framework of mechanistic codon models, each model in the framework hold or relax some of the mentioned simplifying assumptions. Holding or relaxing the three simplistic assumptions results in total to eight different mechanistic models in the framework. Through several experiments on biological datasets and simulations we show that the three simplistic assumptions are unrealistic for most of the biological datasets and relaxing these assumptions lead to accurate estimation of evolutionary parameters such as selection pressure.

## Introduction

The mechanistic codon models have the advantage of having biologically relevant parameters but they suffer from side effect of the simplified assumptions. The reason to hold these assumptions raises from the need to reduce the parameter space of the codon models. However, in considerable amount of data sets these assumptions are not realistic and cause poor results. We particularly consider the following three assumptions:

### Assumption 1: Mutation is homogenous at the nucleotide level

The first simplified assumption made by almost all current mechanistic codon models is that substitution process that is applied to the nucleotide level is homogenous. As a result, these codon models ignore substitution rate heterogeneity across nucleotides and assume one averaged mutation rate matrix for all nucleotide sites. Many studies show that this is an over-simplified assumption and the rate variations across adjacent nucleotides has been observed [1–6].

There are at least two main biological reasons for substitution rate heterogeneity among nucleotide sites namely; selection pressure and mutation bias. Often, the current codon models of evolution take into account the effect of selection pressure by means of *ω* parameter. However, the effect of the mutation bias at the nucleotide level is ignored in most of the mechanistic codon models with rare exceptions such as CNF substitution model [7]. Some studies show that the mutation bias at the nucleotide level should not be neglected. For example in some sequences, there can be more mutation in G and C sites or near insertions and deletions [8, 9]. Furthermore, it has been shown that ignoring substitution rate heterogeneity among nucleotides results to inaccuracy of parameter estimation such as underestimation of branch lengths and transition to transversion rate [2].

### Assumption 2: Double and triple substitutions are neglectable

An important assumption accepted by almost all of the mechanistic codon models, i.e. [7, 10–15] (with rare exceptions such as the SDT model [16] and GPP [17]), is that only a single nucleotide substitution per codon is allowed within a small time interval. Consequently, double and triple substitutions within a codon are ignored and the models that apply this assumption, simply force the corresponding parameters of double and triple substitutions to be set to zero [18, 19]. This results in simpler models and reduces the number of codon transition rate parameters significantly. In particular, the number of paramaters reduces from 1,830 to about 500 that results in faster algorithms.

However, this simplified assumption might be unrealistic as multiple instantaneous substitutions have been observed in real DNA sequences, and mathematical models that consider doubles and triples nucleotide site substitution show more accurate results [16, 20, 21]. For instance, the best estimates of protein evolution have nonzero instantaneous rates of change between amino acids whose codons differ by more than one nucleotide [22, 23]. Moreover, evidences for high frequency of simultaneous double nucleotide substitutions are discussed in [24, 25]. Furthermore, highly frequent small scale genomic events, such as codon bias, gene conversions and sequence inversion can make the double and triple substitutions very likely [16, 26–28]. Although the rates of double and triple substitutions have been estimated to be 2 to 3 orders of magnitude lower than single substitutions [16, 29, 30], studies based on empirical data show that in average, about 25% of codon transitions are double and triple. This further indicates that double and triple substitutions are not neglectable [20, 21].

The only fully parametric model that considers multiple instantaneous substitutions was developed by Whelan and Goldman [16]. It calculates substitution rate matrices for single-, double- and triple-nucleotide mutation separately using the equilibrium frequency of mutated nucleotides and a transition to transversion rate. The three matrices are then combined to calculate the general rate matrix of codons. Evaluation of this model on a large amount of coding sequences showed that this assumption improved the likelihood performance compared to the existing mechanistic models [16].

### Assumption 3: HKY model is good enough at the nucleotide level

The third assumption made in most of the mechanistic codon models including M0 model and its extensions is estimating the transition to transversion rates at the nucleotide level by HKY model [18]. HKY model has only one transition to transversion rate parameter that represents the mutation substitution rate at the nucleotide level. However, more general and sophisticate models can be exploited at the nucleotide level. The research shows that, in most sequences, GTR model is explaining the real nucleotide sequences better than HKY model [31]. Therefore, using GTR model instead of HKY model in codon models may explain the data sets more accurately. But, despite of this fact, the HKY model is applied in many mechanistic models to simplify the model and overcome computational costs.

The need for the above three simplified assumptions raises from reducing the computational complexity of the parameter estimation of the mechanistic codon models. Transition rates of codons and selection pressure are two important parameters of codon models. Next, we explain these two classes of parameters and mention why the three assumptions makes their estimation computationally tractable.

### Transition rates of codons

The first class of biologically plausible parameters in codon based models are transition rates between codons in the codon substitution rate matrix. There are 4,032 potential variables, i.e. 64 × 64 minus 64 diagonal entries, for all possible codon transition rates. Assuming that the transition rates between codons are symmetric and also transition rates to and from stop codons are zero, one can reduce the number of possible variables to 1, 830. This is still a very large set of variables that makes the full estimation computationally demanding. Furthermore, in practice, the input sequences often contains much less information that makes many of these variables dependent and redundant.

Mechanistic-empirical models, such as ECM model, estimate transition rates empirically from a large database and name it exchangeability rate. Then, this matrix is combined with equilibrium frequency of codons to build the codon substitution rate matrix, and is used as a fixed component and is never optimized. Accordingly, one can say, the original ECM has no parameter for estimating sequence-specific transition rates as it assumes that the transition rate matrix is unique across all data sets. This unique transition matrix can be interpreted as an average transition rate matrix which is shown to be effective in practice.

There has been actually two drawbacks for exploiting a precomputed and fixed transition matrix for all sequences. Intuitively, it is expected that the empirical exchangeability matrix perform the best in some data sets, however as the sequences diverges from the data sets used to estimate the rates, the performance of using a fixed empirical exchangeability rate drops down. The second issue with the empirical exchangeability rates is that there is no straight forward way to separate the information of selection pressure and transition to transversion rate of nucleotides from exchangeability rates of codons.

Alternatively, to have computational efficient models and overcome huge information redundancy, mechanistic models accept all or some of the above simplified assumptions and as a result, estimate the large set of codon transition rates by few number of parameters. For example, M0 and its extensions assume all of these variables can be estimated by only one transition to transversion rate parameter, i.e. *κ*, borrowed from HKY nucleotide model.

### Selection pressure

The second class of biological parameter in codon models is selection pressure. The goal is to detect how much perturbation exist in the proportion of nonsynonymous and synonymous substitutions in a sequence compare to the case that the sequence is under neutral evolution. The selection pressure is measured by estimating the rate of nonsynonymous substitutions to synonymous substitutions under selection pressure divided by the same measure under neutral evolution [10]. In order to differentiate between neutral evolution and the alternative selection, codon models consider at least one parameter *ω* multiplied by all nonsynonymous codon transition rates under the alternative selection.

So far, most of the methods that has been developed to estimate selection pressure at the protein level accept all or some of the above mentioned assumptions. The reason behind rate variation across nucleotide sites in a protein coding region is usually assumed to be due to selection pressure and this is modeled by the *ω* parameter [18] under selection, and under neutral evolution the three codon positions have same nucleotide transition rate matrix. Furthermore, the general view is that double and triple substitution rates are not possible under neutral evolution. As it was discussed above, there are biological processes that support relaxing the above mentioned assumptions and holding these assumptions may result to inaccurate estimation of selection pressure in some sequences. Despite the existence of the biological processes that reject these assumptions, the selection pressure estimated under above assumptions is considered as the standard estimation of the selection pressure and most of the mechanistic codon models consider these assumptions for estimating of the selection pressure. Next, we will briefly overview some of the main methods developed to estimate selection pressure rate averaging over all sites and branches.

Counting method was the first approach developed to estimate selection pressure between sequences [18, 32]. It has three main steps, in the first step the synonymous and nonsynonymous sites were counted. Then the synonymous to nonsynonymous differences were counted and in the last step the proportion of differences were calculated and corrected for multiple hits. In the first two steps, in order to decrease the computational complexity, the method holds the second assumption and ignore double and triple substitutions. Moreover in the last step, they do correction for multiple hits using a single nucleotide substitution rate matrix of JC69 model [33]. This implicitly means that they assume mutation homogeneity by using one transition rate matrices for the three codon positions, and furthermore the nucleotide model used in this approach is even simpler than HKY model. Later on, transition-transversion rate and unequal codon frequency were applied to this method [34].

M0 model, a reference mechanistic codon model, holds all three simplified assumptions, i.e. mutation homogeneity, ignorance of double and triple substitution and HKY model at the nucleotide level. As a result its estimation of *ω* is similar to the selection pressure estimated by counting method. The SDT model was the first mechanistic model that relaxed the second assumption and allowed double and triple substitution [16]. Two selection pressure parameters are estimated in SDT model, *ω*_1_ and *ω*_2_. The former is used to show the effect of selection pressure on two neighboring amino acids. However, *ω*_1_ estimates the effect of the selection pressure on codons. It has been shown that *ω*_1_ under-estimate the *ω* parameter estimated by models that ignore double and triple substitution. This clearly shows that if a data set has adequate amount of double and triple substitutions, then the models that ignore these substitutions do not estimate the selection pressure accurately [16]. In the ECM model, by allowing double and triple substitutions and empirical estimation of transition rates, the three above unrealistic assumptions are relaxed for estimation of selection pressure [21]. However, their direct estimation of *ω* is not accurate because the empirically estimated exchangeability rates of codons is also affected by the selection pressure. Accordingly, two different methods have been developed to extract the information about selection pressure from exchangeability rates. In both methods, the goal is to estimate the selection pressure similar to the standard selection pressure, i.e the selection pressure estimated by M0 model.

The first approach, introduced in [21], the rate of nonsynonymous to synonymous substitutions under selection is estimated and divided by a constant rate obtained from *α* – *β*globin gene as the universal rate of nonsynonymous to synonymous substitutions under neutral evolution by means of counting method [32]. As a result to calculate this constant rate, the three above simplified assumptions are considered.

The second method estimates the rate of nonsynonymous to synonymous substitutions under neutral evolution directly from the substitution rate matrix of the ECM model, instead of using the above mentioned constant rate [35]. However, they constraint their approach to hold the three above simplified assumptions. They ignore all double and triple substitution rates in the estimation of nonsynonymous to synonymous substitutions under neutral evolution. Moreover, for the single substitutions they estimate nonsynonymous to synonymous rate ratios for transitions and transversions separately and then average them. This implicitly means that they assume one nucleotide transition to transversion rate for the three codon positions. Furthermore, they assume one transition and one transversion rate over all three nucleotide position similar to HKY model.

It is important to mention that although selection pressure estimation can be lineage or site specific [11, 36–39], in this paper, we focus on the average selection pressure over all lineages and sites.

In this paper, we introduce a framework of a family of models, i.e. *KCM-based model family*, developed by means of relaxing or holding three main assumptions of the current mechanistic codon models. We explained in detail in [40], the most general model of *KCM-based model family* framework, i.e. *KCM_19x_* [40]. We showed that *KCM_19x_* outperformed the currently most used codon models, i.e. M0 and MEC models, with respect to the *AICc*s metric. Furthermore, the biological meaning of the parameters of the *KCM_19x_* were discussed in details through several experiments. This paper focuses on the underlying assumptions in the current mechanistic codon models and presents how much these assumptions are realistic and supported by empirical data sets. The *KCM-based model family* gives an appropriate framework for this purpose.

*KCM-based model family* framework gives us the power to start from the most restricted model, *KCM_2xM0_*, which is proved to be equivalent to the M0 model, and move toward the more complex models. Finally, on the other side of the spectrum, we get to the most general model by relaxing all three restricting assumptions as shown in figure 1. The whole journey is reinforced by a set of simulations.

**Figure 1.**
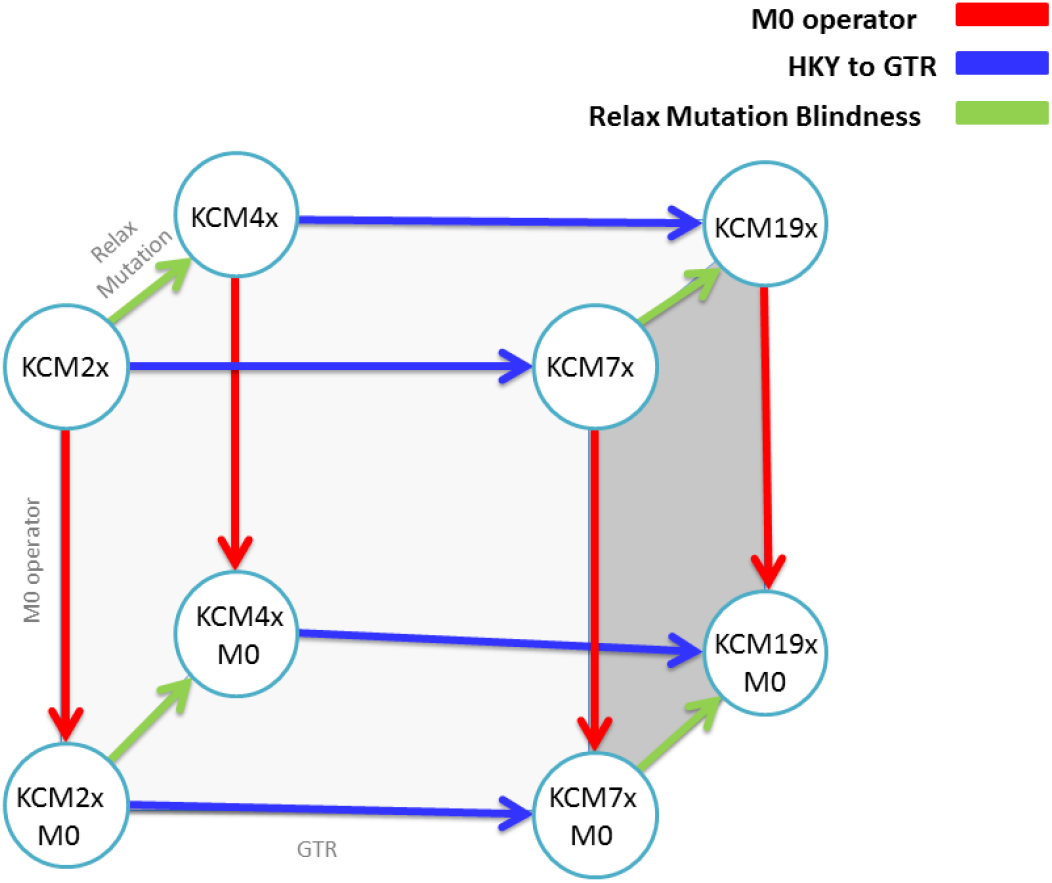
This figure shows the relationships among various models of *KCM-based model family*.

Furthermore, exploiting the same idea, we show statistically what percentage of empirical data sets support each model and its underlying assumptions. As a result, we suggest to use the *KCM-based model family* framework as a tool to classify the empirical data sets and select the model that fits best with the salient characteristics of the data set. Finally, the relationships among parameters of the various models of *KCM-based model family* framework is clarified.

## Materials and Methods

### *KCM-based model family* Framework

In this section, we present the structure and building blocks of models of *KCM-based model family* framework. Generally in models of *KCM-based model family* framework, codons are coded by 3 consecutive nucleotides that are free to vary and our approach to generalize the model is to assume that substitutions occurring within a codon are independent. The nucleotides present in the 3 codon positions can therefore change independently and instantaneously. Each nucleotide *i* within a codon is further modeled by a symmetric substitution matrix *q_i_* similar to GTR or HKY substitution matrices. The models of *KCM-based model family* framework are obtained by combining the 3 matrices at each codon position using Kronecker product.

### *KCM_19x_*, The Generalized Model

To describe the *KCM-based model family* framework, first, we remind the main and the most general model of this family, *KCM_19x_* model [40], that relaxes all three simplified assumptions. Next, we discuss other variations of *KCM-based model family* framework that are constructed by holding some of the mentioned restricting assumptions.

In *KCM_19x_*, each nucleotide *i* within a codon is further modeled by a symmetric substitution matrix *q_i_* that allows distinct rates for each type of substitutions. This is in essence similar to the *GTR* substitution matrix [41], although the state frequencies are not included here:

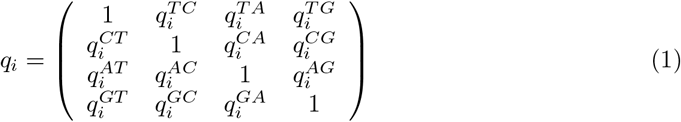

where the rate of change between nucleotides *j* and *k* in the ith codon position is given by 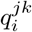 and 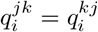. Note that each of the 3 matrices is not normalized at this stage and the 18 parameters (i.e. 6 per nucleotide positions) are free to vary.

The *KCM_19x_* model is obtained by combining the 3 matrices at each codon position using Kronecker product. The result of Kronecker product of the 3 consecutive 4 × 4 matrices at each codon position is a 64 × 64 matrix (hereafter referred to as Kronecker matrix), that represents, after some post-processing described below, the rate transitions between any codons based on the underlying substitution rates of the nucleotides. The initial matrix includes substitutions from and to the 3 stop codons which should be removed. We obtain a 61 × 61 Kronecker matrix for sense codons by removing the 3 rows and columns representing substitutions from and to the 3 stop codons. The Kronecker matrix is then multiplied by the diagonal matrix of the equilibrium frequencies of codons Π:

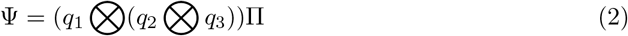

The substitution matrix that is finally representing how codons are changing through time is as follows:

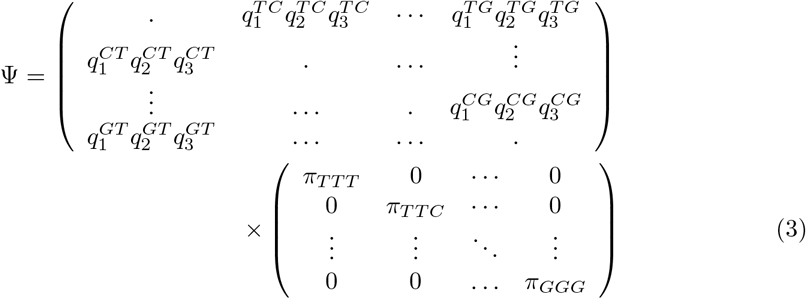

where Ψ is a 61 × 61 matrix, 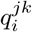 are substitution rates of nucleotide *j* to *k* in the ith position of the codon and *π_m_* stands for equilibrium frequency of codon *m*. The frequencies *π_m_* can be any type of codon frequencies and can be estimated empirically from the data as usually done for codon models [18].

The *KCM_19x_* model can furthermore be extended to include the effect of selection [10]. For every codon *i* and *j*, the parameter *ω*, that represents the ratio between synonymous and nonsynonymous substitutions, is introduced whenever the transition changes the amino-acid coded by the codons, leading to the final matrix *Q*:

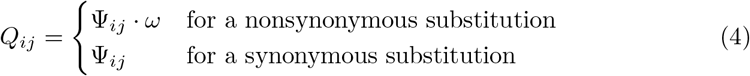

Finally, diagonal elements of the *Q* matrix are fixed to ensure that the row sums of *Q* equal zero, and *Q* is scaled to obtain an average rate of substitution at equilibrium equals to 1. We therefore count double and triple substitutions as single events, that imposes that the branch lengths are measured in expected numbers of substitutions per codon.

#### *KCM_19x_* and the three assumptions

As mentioned before, *KCM_19x_* is the most general model of the *KCM-based model family* framework and it relaxes all three simplified assumptions mentioned in the Introduction section.

In brief, to cover the cases where mutation homogeneity do not hold, it is required to exploit at least three different nucleotide matrices in the model corresponding to the three nucleotide positions in a codon. Although it doesn’t allow a fully free substitution variation among nucleotides as done in nucleotide models mostly with gamma distribution [42–44], it gives three classes of rates variation among nucleotides within codons. Accordingly, we allowed some models of *KCM-based model family* framework including *KCM_19x_* to consider three nucleotide matrices averaged over all first, second and third codon positions, respectively. Furthermore, to relax the second assumption, *KCM_19x_* is developed in a way to allow double and triple substitutions and cope with the dramatic increase of the transition rate parameter space from 528 to 1830. Moreover, we relaxed the third assumption in *KCM_19x_* and various members of *KCM-based model family* by considering GTR model instead of HKY model in the nucleotides level.

#### *KCM_19x_* and selection pressure

In order to estimate the rate of nonsynonymous to synonymous substitutions under selection, in *KCM-based model family* framework including *KCM_19x_*, we use *ω* parameter, in a similar way to M0, ECM and MEC models. However, *KCM_19x_* relaxes the three above assumptions for estimating the rate of nonsynonymous to synonymous substitutions under neutral evolution, and as a result, sometimes it estimate different selection pressure. We developed a method to restrict the estimation of the rate of nonsynonymous to synonymous substitutions under neutral evolution to the simplified assumptions and as a result we get similar estimation as the selection pressure estimated by M0 model, i.e. standard selection pressure. Though, we believe that if in model selection comparison, *KCM_19x_* model that relaxes the above three assumptions fits the data sets better compare to other models, then one should be cautious about the accuracy of the standard selection pressure.

### Extended models of *KCM-based model family* Framework

The general model of *KCM-based model family* framework, i.e. *KCM_19x_*, relaxes the three assumptions mentioned earlier in the Introduction section. By exploiting three 4 × 4 matrices, the general model assumes that mutation acts differently in each nucleotide position of the codon. By definition, it allows double and triple nucleotide substitutions. Furthermore it uses GTR substitution matrix for nucleotide building blocks.

Accordingly, there can be eight combination of the assumptions and for each combination there can be a tailored model in *KCM-based model family* framework. the models are obtained by reducing the generalized model, i.e.*KCM_19x_*, holding the corresponding combination of assumptions. Following, the variations are presented and discussed.

#### *KCM-based model family*, double and triple substitution rates

In order to restrict *KCM_19x_* to a model that allows only single codon substitutions, we introduced the *M*0 operator *α* ∈ {0,1} as follows:

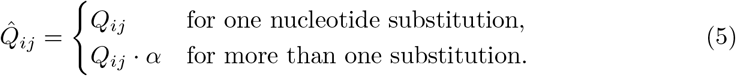

With *α* = 0, double and triple substitutions are not allowed (*KCM* M0). The variation of *KCM_19x_* that doesn’t allow double and triple substitution is named *KCM_19xM0_*.

#### *KCM-based model family* and homogenous mutation rate

The mutation homogeneity assumption can be obtained by reducing the three nucleotide matrices of *KCM_19x_* to one 4 × 4 matrix that is kronecker multiplied by itself three times. This variation called *KCM_7x_*.

Intuitively, one can obtain both mentioned assumptions simultaneously by restricting *KCM_19x_* by applying M0 operator, i.e. ignoring double and triple substitutions, and reducing three nucleotide matrices to one. We call this version that holds both assumptions, *KCM_7xM0_*.

#### *KCM-based model family*, HKY vs. GTR

In addition to the above variations, one can change the pattern of 4 × 4 building block matrices from GTR to HKY as follow and produces other extensions of *KCM_19x_*.

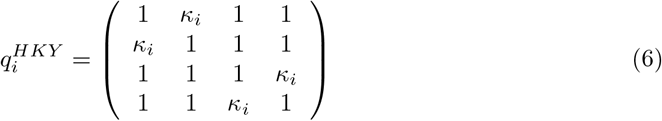

where *κ_i_* is the parameter responsible for any transition-transversion change in the *i*th nucleotide position of the codon. This change reduces number of parameters of each 4 × 4 building block matrices from 6 to 1. Relaxing or restricting the two above assumptions on the models that use HKY model at the nucleotide level results in four more variations of *KCM_19x_*.

In particular, holding HKY assumption and relaxing the other two main assumptions results in a model that we name it *KCM_4x_* that uses three different 4 × 4 HKY building block matrices. Considering mutation homogeneity results in a model that can be called *KCM_2x_* with one 4 × 4 HKY matrix, Kronecker multiplied by itself three times. Finally, ignoring double and triple substitutions results in *KCM_4xM0_* and *KCM_2xM0_* by means of applying M0 operator on *KCM_4x_* and *KCM_2x_*, respectively. As we present in the following theorem, it is mathematically straight forward that *KCM_2xM0_* is equivalent to M0 model.

##### Theorem 1.

*Let* KCM_2xM0_ *be the most restrictive model of* KCM-based model family *framework with one nucleotide matrix inspired from HKY model. Then* KCM_2xM0_ ≡ *M*0.

*Proof*. Assuming HKY matrix as the nucleotide building blocks of *KCM_2xM0_* model, then the substitution matrix of codons is constructed as follow:

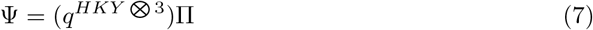

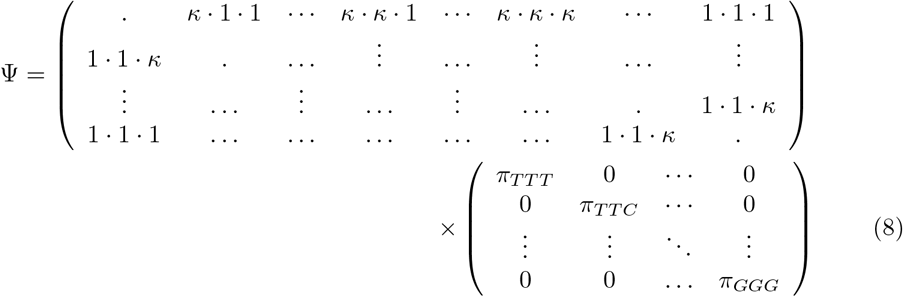

Since the model does not allow double and triple substitutions, M0 operator is applied. Therefore, in the above Kronecker matrix all double and triple substitutions are replaced with 0,

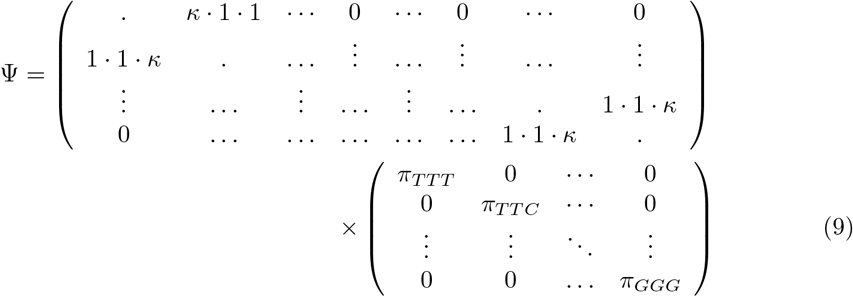

The above matrix multiplication is the matrix representation of M0 model formula proposed for calculating codon substitution matrix.

Therefore M0 model is a subset of *KCM-based model family* framework and a nested model of *KCM* M0 family. Hereafter M0 and *KCM_2xM0_* are interchangeably used.

It also worth to mention explicitly that *KCM_4xM0_* and *KCM_4x_* estimate the three different parameters of transition-transversion rates for each nucleotide position by means of the three different 4 × 4 HKY matrices. *KCM_2xM0_* and *KCM_2x_* estimate one transition-transversion rate parameter for all transition-transversion substitutions in codon rate matrix.

In summary, *M*0 operator, number of 4 × 4 nucleotide matrices and number of parameters in each 4 × 4 nucleotide matrix are three gadgets that generate 8 different variations of *KCM-based model family*. These models can cover all combinations of the three mentioned assumptions. Table 1 shows a summary of different variations of *KCM-based model family*, their formulation, number of parameters and assumptions they consider.

**Table 1.** Different extensions of the *KCM-based model family*. The symbol *q_i_* refers to the type of substitution matrix at each nucleotide position.

| Model description | No. params | mutation homogeneity | Double, Triple Neglected | HKY vs GTR |
| --- | --- | --- | --- | --- |
| $M0 \equiv KCM_{2xM0} \quad \Gamma = (q_1^{HKY})^{\otimes 3}$ | 2 | Hold | Hold | HKY |
| $KCM_{4xM0} \quad \Gamma = (q_1^{HKY} \otimes q_2^{HKY}) \otimes q_3^{HKY}$ | 4 | Relax | Hold | HKY |
| $KCM_{7xM0} \quad \Gamma = q_1^{\otimes 3}$ | 7 | Hold | Hold | GTR |
| $KCM_{19xM0} \quad \Gamma = (q_1 \otimes q_2) \otimes q_3$ | 19 | Relax | Hold | GTR |
| $KCM_{2x} \quad \Gamma = (q_1^{HKY})^{\otimes 3}$ | 2 | Hold | Relax | HKY |
| $KCM_{4x} \quad \Gamma = (q_1^{HKY} \otimes q_2^{HKY}) \otimes q_3^{HKY}$ | 4 | Relax | Relax | HKY |
| $KCM_{7x} \quad \Gamma = q_1^{\otimes 3}$ | 7 | Hold | Relax | GTR |
| $KCM_{19x} \quad \Gamma = (q_1 \otimes q_2) \otimes q_3$ | 19 | Relax | Relax | GTR |

In table 1 one can see classification of the variations *KCM-based model family* and M0 based on the mentioned underlying assumptions. Having a model family has two advantages, (1) one can determine the effectiveness of the assumptions about a data set at hand by comparing the performance of each model and (2) the model appropriate for the data set can be applied to the data set for accurate estimation of parameters.

The illustrative relationships among various models of *KCM-based model family* are presented in figure 1. Starting from the simplest model of the framework, i.e. *KCM_2xM0_* (M0 model), one can find a path to any other model by holding or relaxing the three assumptions. For example, starting from M0 model, i.e. *KCM_2xM0_*, by relaxing mutation homogeneity assumption we can reach to *KCM_4xM0_*. On the other side, by changing HKY model at the nucleotide level to GTR model, the model moves toward the *KCM_7xM0_*. Moreover, by applying M0-operator the model moves toward the *KCM_2x_*.

## Results

In this section, we will verify our claims through a wide range of experiments and present the intuitive role of parameters in the models of *KCM-based model family* framework.

### Empirical data sets

#### Model selection

We applied the different variants of *KCM-based model family* models to the empirical data sets, and we estimated the Δ*AICc* of any model with respect to the other models in the framework. Table 2 shows what percentage of the data sets support the superiority of each model over the other models. The last row shows the percentage of the data sets that the proposed model outperforms all other models with 99% probability. Furthermore, table 6 shows the best model for each data set with the detail of the 70 empirical data sets.

**Table 2.**
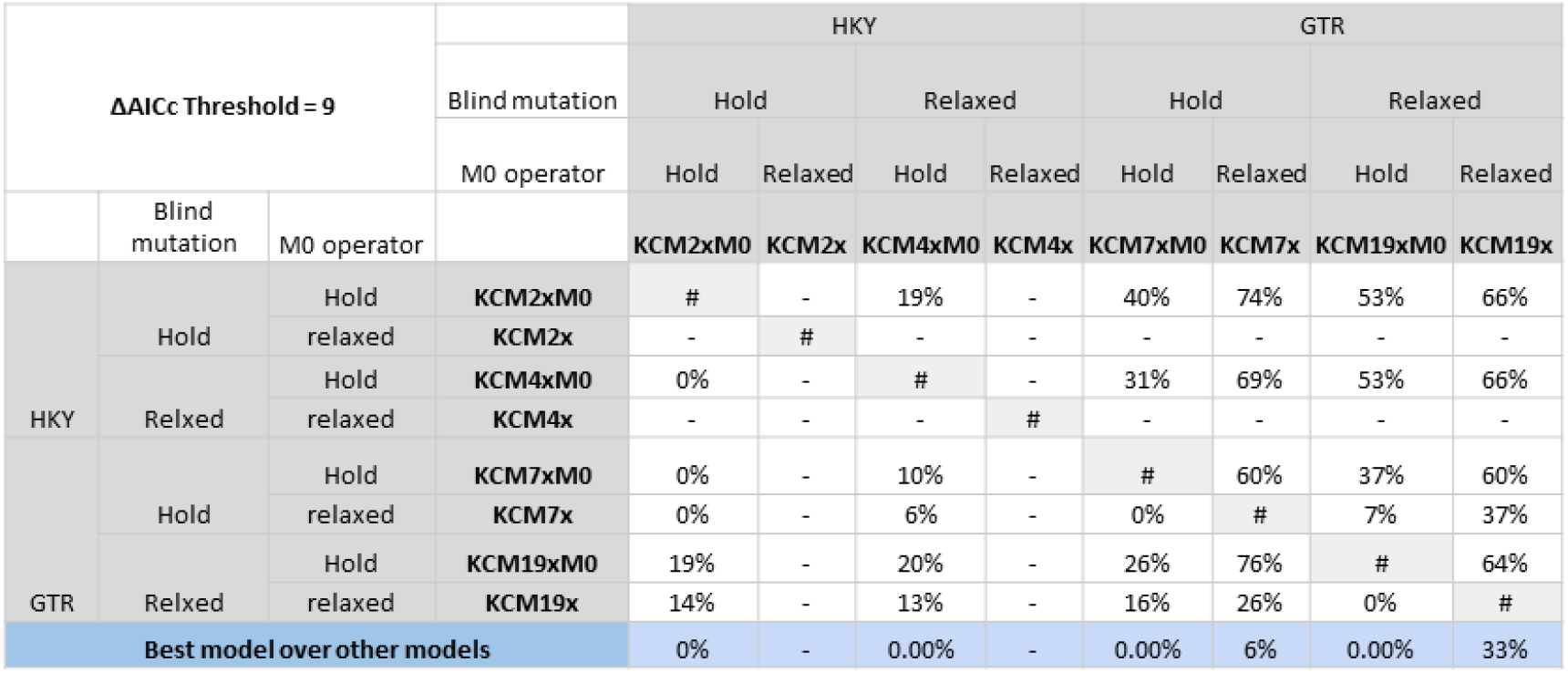
This table shows the percentage of the empirical data sets, selected from the Selectome database, in favor of each model. In the last row, the percentage of the data sets that a model is superior over all other models are shown.

Comparing the most general model, *KCM_19x_*, with the simplest model *KCM_2xM0_* (M0 model). There are about 10% of data sets in favor of the simplest model. Although the main three assumptions of mutation homogeneity, ignorance of double and triple substitutions and GTR vs HKY are imposing strong constraints on the model of evolution, there are some data sets that seem to fit this simple model. We conclude that mutation heterogeneity and double and triple nucleotide substitution within codon is very low in these data sets. Furthermore, the transition rates between nucleotides can be taken into account by one parameter.

Comparing *KCM_19x_* with *KCM_7x_* shows what percentage of data sets hold mutation homogeneity while both models allow double and triple substitutions. For about 26% of data sets the best model is *KCM_7x_* with 99% probability relative to *KCM_19x_*, and for 37% of data sets *KCM_19x_* is the best model.

*KCM_19x_* and *KCM_19xM0_* both relax the mutation homogeneity assumption. However, *KCM_19x_* allows double and triple substitutions. The results show that allowing only single substitution assumption is not a good assumption for any data sets.

There are 26% of data sets in favor of *KCM_7xM0_* compared to *KCM_19xM0_*. This shows that when we do not allow double and triple substitutions 26% of data sets hold mutation homogeneity assumption and rate variation across nucleotides within codons can be ignored. As mentioned above, comparing *KCM_7x_* and *KCM_19x_*, allowing double and triple substitutions does not change this percentage. Meaning that mutation homogeneity is totally independent from holding or relaxing double-triple substitution assumption. Moreover, comparing *KCM_7xM0_* with *KCM_19x_* shows 16% of data sets hold both main assumptions of mutation homogeneity and negligible rate of double-triple substitution rates under GTR model.

Comparing *KCM_2xM0_* with *KCM_7xM0_* shows the percentage of data sets preferring HKY model of nucleotides over GTR model when we analyze codons. There is no data set in favor of HKY model in the nucleotide level.

No data set is in favor of *KCM_2xM0_* compare to *KCM_4xM0_*. Comparing *KCM_2x_* and *KCM_4x_* to *KCM_2xM0_* shows that all data sets are in favor of *KCM_2xM0_*, because by allowing double and triple substitutions the number of variables that a model should estimate increase dramatically, however the number of parameters of the model does not increase enough. Therefore in order to allow double and triple substitutions one needs at least a GTR model (six parameters) for the 4 × 4 nucleotide matrix.

Fig 9 shows the pie chart of the Δ*AICc* of the two mentioned models over 70 mentioned sample data sets. In 60% of data sets, *KCM_7x_* fits better than *KCM_7xM0_* and the fit is significant, i.e. the Δ*AICc* > 9. It can be concluded that neglecting double and triple substitutions is not realistic in more than 60% of the targeted data sets. For 11% of data sets, we can not say which model is the best model with high confidence, i.e. 9 > Δ*AICc* > 4. For 29% of data sets, there are substantial evidence for alternative model, *KCM_7xM0_*. However there is no data set that *KCM_7xM0_* model fits significantly better than *KCM_7x_* model.

Fig 10 shows the Δ*AICc* of *KCM_7x_* versus *KCM_19x_*. The results shows that *KCM_7x_* is a good model in terms of fitting to the data sets. For example, as shown in table 2 in 74% of data sets this model fits better than *KCM_2xM0_*, but *KCM_2xM0_* can never significantly fit better compare to *KCM_7x_*. Having this in mind, one can see in figure 10, holding double and triple substitution assumption, mutation homogeneity assumption is unrealistic for 37% of data sets. However, for 26% of data sets mutation homogeneity is a realistic assumptions, i.e. *KCM_7x_* fits better compare to *KCM_19x_*. 21% of data sets are in inconclusive region, and for 16% the simpler model is good enough, i.e. *KCM_7x_*. The same results are obtained comparing *KCM_7xM0_* with *KCM_19xM0_*, meaning that *KCM_19xM0_* and *KCM_7xM0_* models fit better on 37% and 26% of data sets, respectively.

Combining the two observed results, wherever *KCM_7x_* fits better simultaneously over *KCM_7xM0_* and *KCM_19x_*, it can be concluded with higher confidence that mutation homogeneity can be a proper assumption for some data sets while neglecting double and triple substitutions is not. Our experiment shows that in 6% of data sets, *KCM_7x_* fits better simultaneously over *KCM_7xM0_* and *KCM_19x_*. In particular, figure 2 shows that 6% of the data sets (shown with red dots) are in favor of mutation homogeneity but against neglecting double and triple substitutions while 0% of data sets are in favor of opposite assumptions, i.e. taking into account mutation heterogeneity and ignoring double and triple substitutions. Furthermore, 31% of the data sets (shown in green dots) prefer relaxing mutation homogeneity assumption but allowing double and triple substitutions. No data set is in favor of mutation homogeneity but neglecting double and triple assumptions. The blue dots shows the set of the data sets that we can not identify the best model for them with high probability.

**Figure 2.**
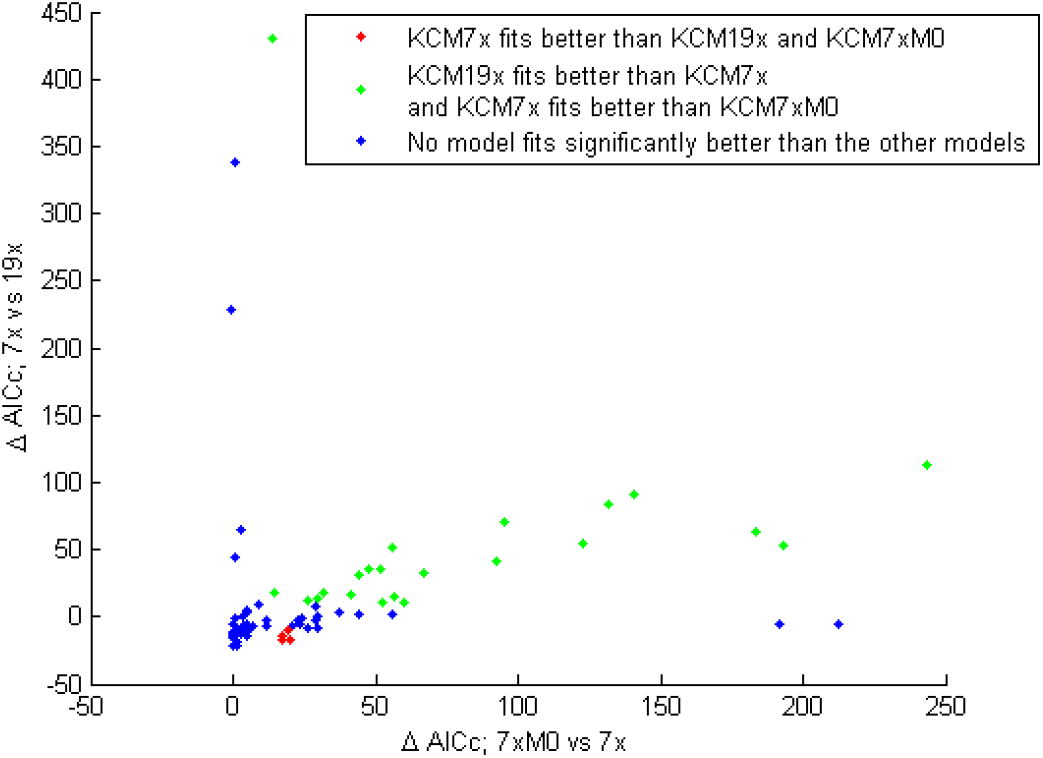
This figure shows a 2 dimensional view of the percentage of empirical data sets that are in favor of *KCM_7x_* model compare to *KCM_19x_* and *KCM_7xM0_* models.

There are 6% of data sets, in our sampled group, where *KCM_7x_* outperforms all other models according to the Δ*AICc* metric. In 33% of data sets *KCM_19x_* outperform all other models. The last row of the table 2 shows the percentage of data sets the proposed model outperforms all other models with 99% probability.

#### Parameter estimation

##### Average transition-transversion rate ratio

The results on empirical data sets also confirms that the *KCM_2xM0_* and M0 models are equivalent. Their parameters, *κ* and *ω*, are estimated exactly the same as well as the likelihood values, i.e. the average ratios of *κ*, *ω* and likelihood estimated by the *KCM_2xM0_* to the corresponding parameters estimated by the M0 model is 1.0000 with the standard deviation equal to 0.

We compare the average transition-transversion rate ratio parameter, *R* [18], of the *KCM_2xM0_* model (hereafter reference *R*) with the *R* estimated by the other models of *KCM-based model family* framework.

The estimated average *R* of the *KCM_4xM0_* model shows good approximation of the reference *R* (fig 11). Moreover, we perform the Wilcoxon rank sum test. The results emphasize that the samples are coming from the same distribution (table 4). However, as is presented in figure 11, there are some outliers. Focusing on the extreme outlier, i.e. data set number 20, the Δ*AICc* comparison shows that no model fits to this data set with high probability. Furthermore, *KCM_7xM0_* is the model with minimum AICc for this data set compare to *KCM_4xM0_* and the other complex models that allow mutation heterogeneity, i.e. *KCM_19x_* and *KCM_19xM0_* models (Table 6). Therefore, we conclude that this data set have homogenous mutation rate. This is maybe due to low level of mutational bias in this data set. Accordingly, *KCMj_xM0_* and the other complex models, i.e. *KCM_19x_* and *KCM_19xM0_* models, may estimate the parameters inaccurately.

Comparing *R* estimated by *KCM_7x_* and *KCM_7xM0_* with the reference *R*, the results show that *KCM_7x_* and *KCM_7xM0_* measure the average transition-transversion ratio very similar to the reference *R* (figure 12, and table 4).

Focusing on the *KCM_19x_* and *KCM_19xM0_* models, the three *R* parameters are estimated for each 4 × 4 nucleotide matrices of *KCM_19x_* and *KCM_19xM0_*. The average of them show good approximation of the reference *R*. (figure 3 without 5 extreme outliers, and table 4). However, *KCM_19x_* and *KCM_19xM0_* do not approximate *R* as accurately as *KCM_2xM0_*, maybe due to the fact that *KCM_19x_* and *KCM_19xM0_* models are mapping a scalar, i.e reference *κ*, to a 19 dimensions parameter space and it may cause some loss of information.

**Figure 3.**
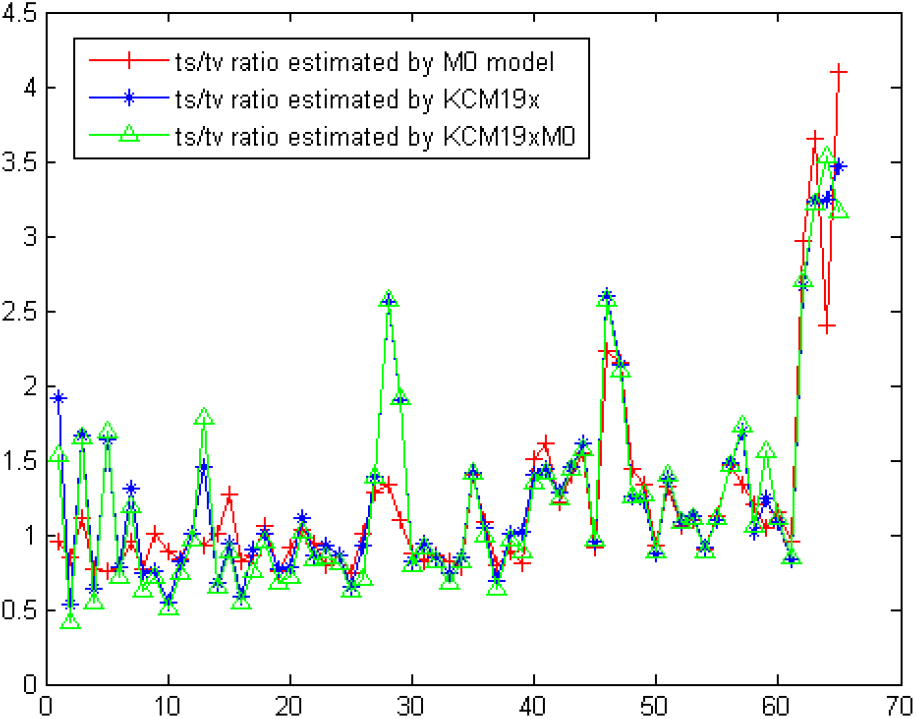
Estimation of the average rate of transition and transversion of *KCM_19x_* and *KCM_19xM0_* models for the empirical data sets selected from the Selectome database. The data sets are sorted based on the *ω* value estimated by *KCM_2xM0_* model.

##### Selection pressure

We investigate the modifications of selection pressure parameter as the underlying assumptions of the model changes. If we consider the current underlying assumptions made by most of the selection estimation methods mentioned in the Introduction section, then we can choose *ω* estimated by the *KCM_2xM0_* model as the reference selection pressure parameter.

Comparing the *ω* parameter estimated by *KCM_4xM0_* and reference *ω*, figure 13 shows they are very similar. We perform the Wilcoxon rank sum test for selection pressure parameters. The result emphasize that the samples are coming from the same distribution (table 4).

The *KCM_7xM0_* estimate *ω* very similar to the reference *ω* (figure 14, table 4). *KCM_7x_* model gives good approximation of *ω* with very similar trend compare to reference *ω* (figure 14, table 4). However, the *ω* estimated by *KCM_7x_* in some cases underestimate the reference *ω* (figure 14).

Comparing *ω* estimated by *KCM_19x_* and *KCM_19xM0_* with the reference *ω*, as shown in figure 4, in most of the cases *KCM_19x_* and *KCM_19xM0_* underestimate *ω*, because their underlying assumptions for estimating selection pressure is different from the underlying assumptions of the reference *ω* as we discussed in the Introduction section. However, the difference is not significant and Wilcoxon rank sum test shows that *ω* estimated by *KCM_19x_* and *KCM_19xM0_* and reference *ω* are coming from the same distribution. (table 4)

**Figure 4.**
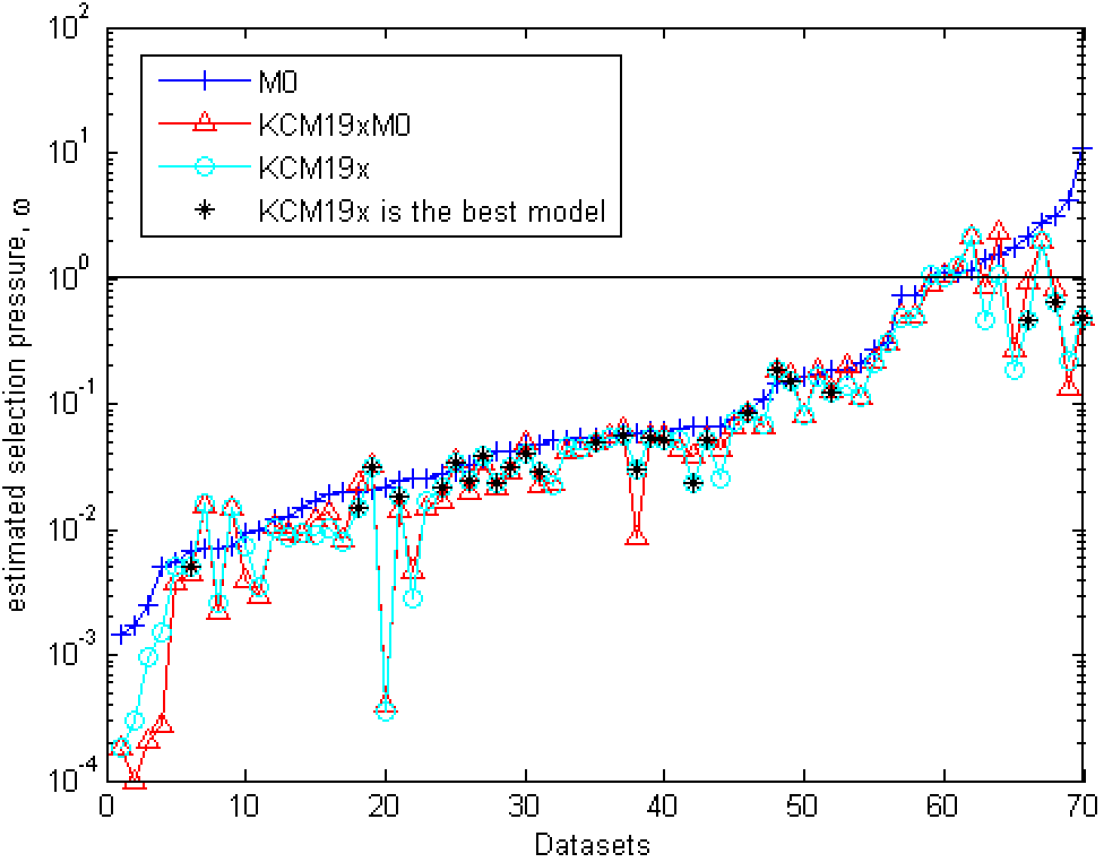
Selection pressure, *ω*, estimated by *KCM_19xM0_*, *KCM_19x_* and *KCM_2xM0_* models for empirical data sets selected from the Selectome database (y-axes is in logarithmic scale).

Comparison of the value of *ω* estimated by *KCM_19xM0_* and *KCM_19x_* to *KCM_2xM0_* shows the effect of relaxing HKY and homogenous mutation assumptions at the nucleotide level under ignoring and allowing double and triple substitution rates. The trend of estimated *ω* in most cases are similar, however as discussed in [40], *ω* estimated by *KCM_19x_* and *KCM_19xM0_* needs correction (*ω_KCM_*) to estimate similar selection pressure estimated by M0 model. Figure 5 shows that *ω_KCM_* has very similar estimation as *ω* estimated under M0 model.

**Figure 5.**
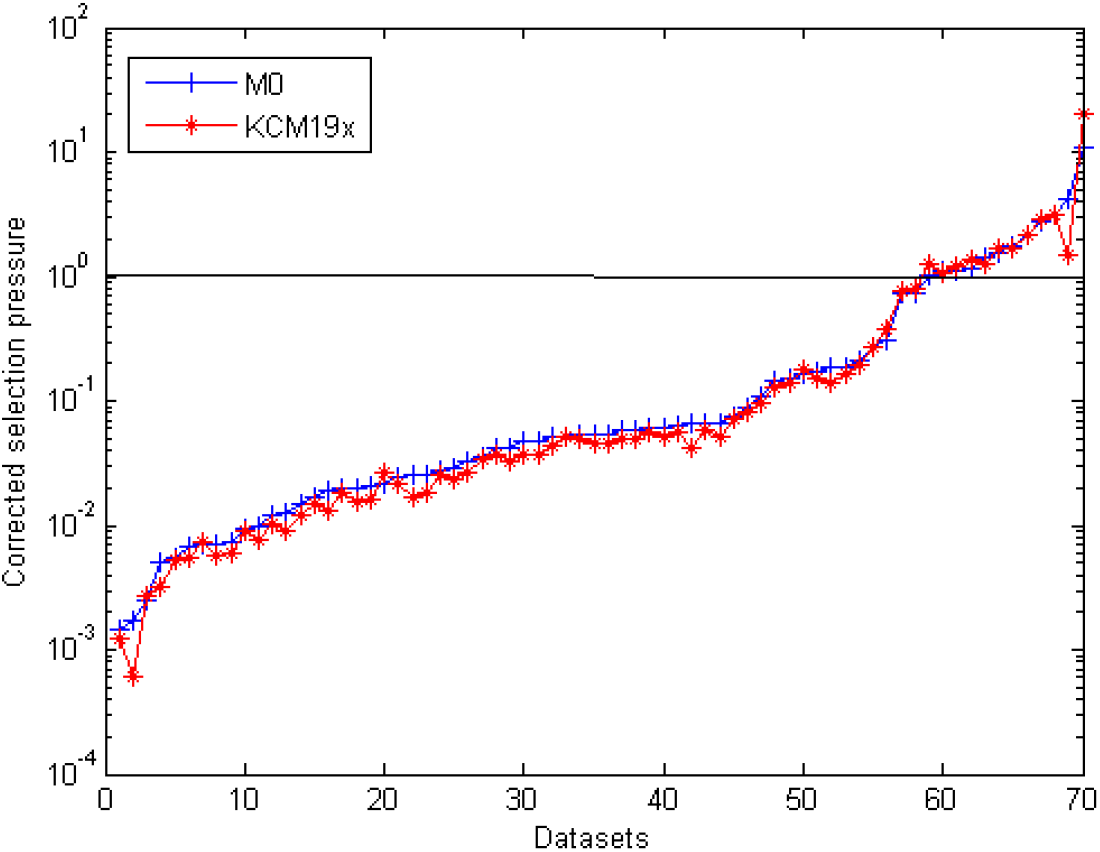
*ω* estimated by *KCM_19x_* (after correction *ω_KCM_*) and *KCM_2xM0_* models. The data sets are sorted based on the *ω* value estimated by *KCM_2xM0_* model (y-axes is in logarithmic scale).

As presented in figure 4, *ω* estimated by *KCM_19x_* without correction in most of the cases have similar trend as the reference *ω*. However, there are few data sets that the estimated *ω* by *KCM_19x_* and M0 models have significant differences. For the data sets that *KCM_19x_* is not chosen as the best model with high probability, we rely on the standard selection pressure. However, as shown in figure 4 there are data sets, such as data set number 70 (figure 4), that *KCM_19x_* is chosen as the best model with high probability and *ω* is significantly different from the reference selection pressure. The fact that *KCM_19x_* is chosen as the best model in these data sets means that these data sets have adequate amount of double and triple substitutions and mutational heterogeneity. Therefore, *KCM_19x_* estimate the parameters more accurately compare to simple models. Accordingly, we suggest that one should be cautious about relying on the estimated selection pressure in such data sets (The details of these datasets are presented in table 6).

### Simulated data sets

In this section, we present the results of model selection on simulation data sets. We perform a set of simulations based on the models of *KCM-based model family* framework. The goal of the simulations is to generate data sets in a way to make three mentioned assumptions gradually weaken or strengthen. This is done either by gradual modification of the 4 × 4 matrices of the models of *KCM-based model family* framework, or by direct modification of the 61 × 61 matrices. In both cases, the gradual modification is done by applying random noise to the entities of the corresponding matrices. In particular, we draw the random noise from a normal distribution justified by the fact that normal distribution approximates the actual distribution of the random noise in the nature reasonably well. The noise is drawn from 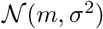, where *m* is the corresponding entity of the matrix and *σ*^2^ ∈ {0.05 * *m*, 0.1 * *m*, 0.15 * *m*, 0.20 * *m*}, i.e. relative noise around the initial value of the entities. We have made sure that the corresponding distributions never cover the negative numbers.

Recalling the cubic illustration of *KCM-based model family* (figure 1), the applied procedure is as following. First, we choose a side on the cube to make the simulations tractable. We would like to study the gradual movements of one model towards the others in the same plain. Second, on each side of the cube, we choose a reference model and we obtain data sets based on the reference model, by simulation. Then, we gradually modify the entities of 4 × 4 matrix of the reference model by applying low level of noise, 5%, i.e. 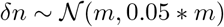. By increasing the level of noise, the simplifying assumptions are gradually violated where the simulated data sets push towards the other models on the same plain. For example on figure 15, we start from 4 × 4 HKY matrix of *KCM_2xM0_*, and the goal is to modify the entities of this matrix with gradual steps and move towards GTR matrix and as a result *KCM_7xM0_* model. After each step of the modification, we generate the data sets related to that level of the noise and store them for the next steps of the study.

It is worth to mention that the simulations are performed only on 2 sides of the cubic illustration (Figure 6). The first side is the lower side containing all models that ignore double and triple substitutions, i.e. the side that is defined by *KCM_2xM0_*, *KCM_4xM0_*, *KCM_7xM0_* and *KCM_19xM0_* as vertices. The second side is the right hand side that is defined by *KCM_7x_*, *KCM_7xM0_*, *KCM_19x_*, *KCM_19xM0_*, i.e. *advanced KCM* models. We exclude the sides that involve *KCM_2x_* and *KCM_4x_*, because these models perform poorly even compare to *KCM_2xM0_* model. The intuitive reason to observe such a poor performance is that when the double and triple substitutions are allowed, the variable space is expanded dramatically from 526 single substitutions to 1830 single, double and triple substitutions. Therefore, in *KCM_2x_* the *κ* parameter should estimate much more transition rates compare to the *κ* of *KCM_2xM0_*. This increase in the variable space reduces the performance of *KCM_2x_* compare to *KCM_2xM0_*. As a result, the likelihood of *KCM_2x_* is always less than the likelihood of *KCM_2xM0_* model. Even though in *KCM_4x_* model, we increase the number of parameters responsible for estimating transition rates to three parameters (*κ*_1_, *κ*_2_, *κ*_3_), the likelihood of *KCM_4x_* is still lower than the likelihood of *KCM_2xM0_*.

**Figure 6.**
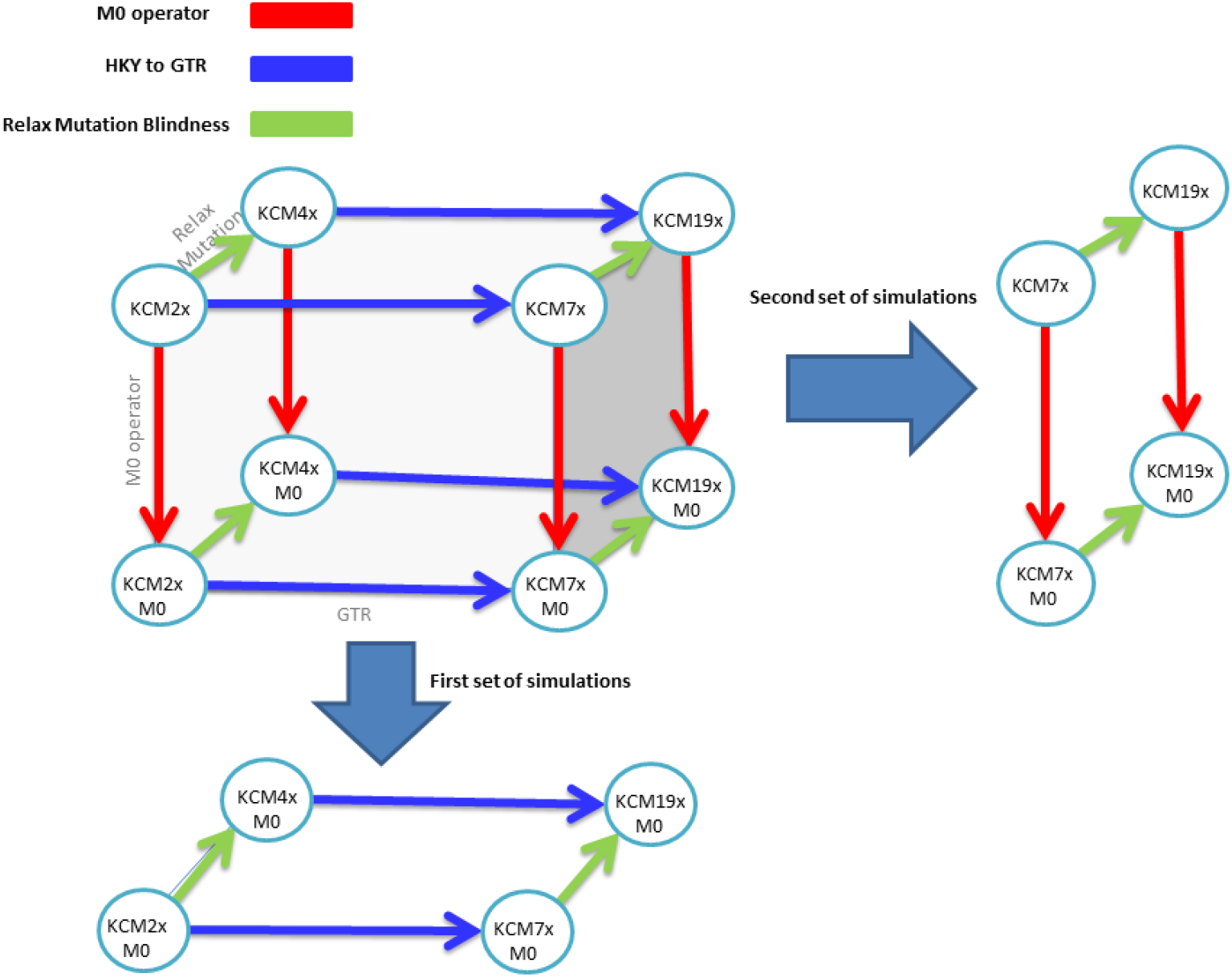
This figure shows the cubic illustration of the models and the two sides that two sets of simulations performed on them.

#### First set of simulations

First, we discuss the first set of simulations, i.e. figure 15. We start from the *KCM_2xM0_* model and we add different types and percentages of noise to move towards *KCM_4xM0_*, *KCM_7xM0_* and *KCM_19xM0_* models. Refer to S1_Simulation1 for more details about this set of simulations.

Figure 7 shows the result of pair by pair model selection between *KCM_2xM0_* and the three other models. *KCM_4xM0_* model has a gradual progress over *KCM_2xM0_*. At 5% type 2 noise, mutation heterogeneity, *KCM_4xM0_* model fits better compare to *KCM_2xM0_* over 2% of data sets, at 10% type 2 noise *KCM_4xM0_* fits better compare to *KCM_2xM0_* model in 14% and at the maximum level of noise, 20%, *KCM_4xM0_* is the best model on 42% of data sets compare to *KCM_2xM0_* with 99% probability.

**Figure 7.**
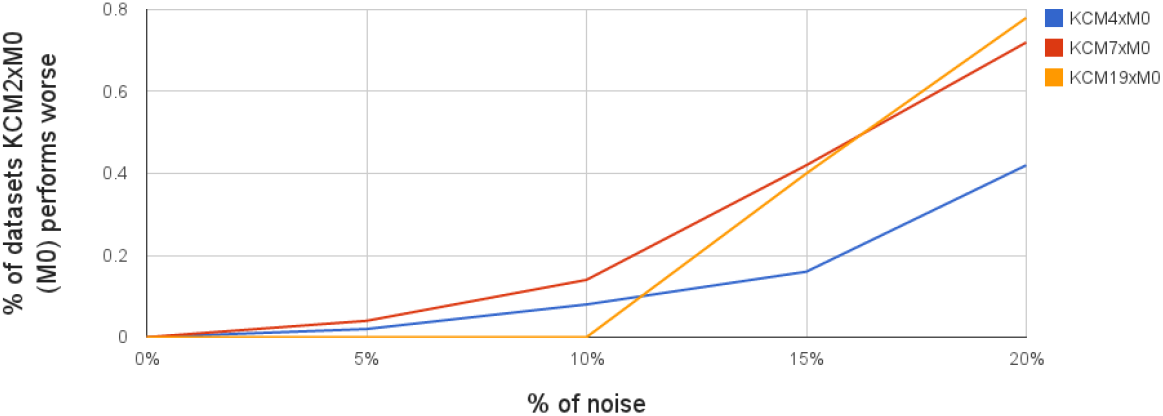
The percentages of data sets that M0 model perform worse than the alternative models on first set of simulations, i.e. M0 space of the cubic illustration.

As Figure 7 presents, comparing *KCM_7xM0_* over *KCM_2xM0_*, at 5% noise, *KCM_7xM0_* fits better compare to *KCM_2xM0_* on 4% of data sets. By increasing the noise to 10%, 15% and 20%,*KCM_7xM0_* fits better compare to *KCM_2xM0_* on 14%, 42% and 72% data sets, respectively (Table 3). The results of comparison between *KCM_19xM0_* and *KCM_2xM0_* shows *KCM_19xM0_* is overfitting the data for 5% and 10% noise level. However, it fits better compare to *KCM_2xM0_* for 15% and 20% level of noise (Table 3).

**Table 3.** Class 1 simulation, percentage of the data sets the corresponding model fits better with respect to *KCM_2xM0_* model, the Δ*AICc* > 9.

| Model | Level of noise |  |  |  |
| --- | --- | --- | --- | --- |
|  | 5% | 10% | 15% | 20% |
| $KCM_{4xM0}$ | 2% | 8% | 16% | 42% |
| $KCM_{7xM0}$ | 4% | 14% | 42% | 72% |
| $KCM_{19xM0}$ | 0 | 0 | 4% | 78% |

Moreover, we investigate the effect of selection pressure on the models that relax mutation homogeneity assumption by starting from *KCM_7xM0_* and moving towards *KCM_19xM0_* with different selection pressures, i.e. *ω* = [0.2,1, 5]. Figure 8 shows the effect of selection pressure on the performance of *KCM_19xM0_* model with respect to *KCM_7xM0_*. Under homogenous mutation assumptions, i.e. 0% noise, *KCM_19xM0_* behaves very similar under different selection pressures. As the type 2 noise increases and mutation heterogeneity is introduced at the nucleotide level, *KCM_19xM0_* fits better compare to *KCM_7xM0_* on very similar percentage of data sets under negative and neutral evolution. However, its behavior is a bit different under positive selection. At 10% noise, *KCM_19xM0_* fits better than *KCM_7xM0_* on twice more data sets with positive selection compare to data sets with purifying and neutral selection. At 15% noise, it fits better than *KCM_7xM0_* on less data sets with positive selection compare to data sets with other types of selection. Finally, at 20% noise, *KCM_19xM0_* fits better than *KCM_7xM0_* on same amount of data sets under the three types of selection.

**Figure 8.**
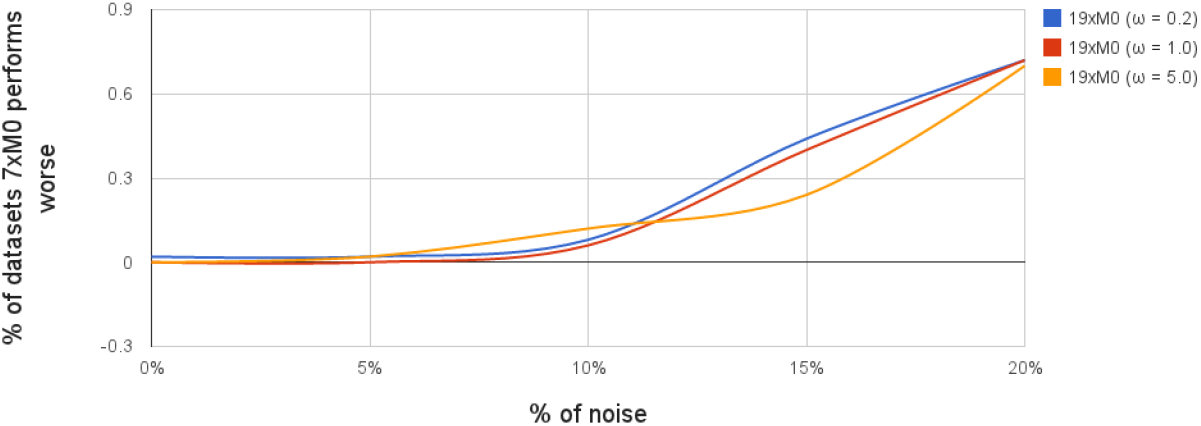
The percentages of data sets that *KCM_7xM0_* model perform worse than the *KCM_19xM0_* model on first set of simulations with three different *ω* values, *ω* ∈ 0.2,1, 5.

#### Second set of simulations

Focusing on the second set of simulations (figure 16)(Refer to S2_Simulation2 for more details about this set of simulations), comparing *KCM_7x_* with *KCM_19x_* model, at 5% and 10% type 2 noise (mutation heterogeneity), there is a clear effect of over-fitting and *KCM_19x_* fits better compare to *KCM_7x_* only on 2% of the data sets. However, as the level of noise increases to 15% and 20%, *KCM_19x_* fits better compare to *KCM_7x_* on 18% and 46% of the data sets, respectively.

Applying type 3 noise, *KCM_7x_* moves toward *KCM_7xM0_*. However, even for the data sets simulated with 0% rate of double and triple substitutions, *KCM_7xM0_* is unable to fit significantly better than *KCM_7x_* for any of the data sets.

Comparing *KCM_19xM0_* model with *KCM_7x_* model, the percentage of the data sets where *KCM_19xM0_* fits significantly better than is 0,0,4%, 36% and 64%, with level of noise 5%, 10%, 15%, 20% respectively. These results are very similar to the percentage of the data sets that *KCM_19xM0_* fits significantly better than *KCM_7xM0_*.

*KCM_19xM0_* never fits significantly over *KCM_19x_* independent from the level of type 3 noise.

## Discussion

In this paper, we investigated the biological relevance of the three mentioned simplified assumptions hold by most of the current mechanistic models. In brief, the simplified assumptions are: i) mutation homogeneity in the nucleotide level, ii) neglecting double and triple substitutions and iii) using HKY model in the nucleotide level. However, some biological processes such as small scale mutations and mutation bias violate these assumptions. Thus, the highly used mechanistic models that hold these assumptions may fail to capture the characteristics of the data sets. To overcome this shortcoming, some of the mechanistic models relax one or two of these assumptions, for example [16] allows double and triple substitution, [7] developed a model that allows mutation heterogeneity and uses GTR in the nucleotide level, and [15] proposed a model that uses GTR at the nucleotide level. However, the idea of allowing double and triple substitution have not been clearly developed in the mechanistic codon models and no study has investigated the effect of the other two assumptions under allowing or ignoring double and triple substitutions.

To address this issue, we proposed a framework of mechanistic codon models, *KCM-based model family* framework, that generates 8 mechanistic codon models by holding or relaxing the assumptions. Having such a framework at hand, the idea simply is to apply different models in *KCM-based model family* framework on a data set and find out the model that performs better than the others. Then, one can associate the corresponding assumptions of the best fitted model to the salient characteristics of the the data set at hand.

Our results show that more than 60% of the empirical data sets are in favor of the models that allow double and triple substitutions. Figure 9 shows the results of comparing *KCM_7xM0_* and *KCM_7x_* on the empirical data sets. *KCM_7xM0_* never outperforms *KCM_7x_* significantly. This shows that the models that ignore double and triple substitutions, i.e. *KCM_19xM0_* and *KCM_7xM0_* perform worst than their alternative models with the same amount of parameters, *KCM_19x_* and *KCM_7x_*, respectively. These results are in agreement with the results that have been mentioned in the literature. For example, it has been shown that double and triple substitutions are the second most important parameters of the codon models after selection pressure parameter, based on Principle Component Analysis (PCA) [45]. Furthermore, the total rate of double and triple substitution estimated by the ECM matrix from empirical data sets is about 25% [21].

**Figure 9.**
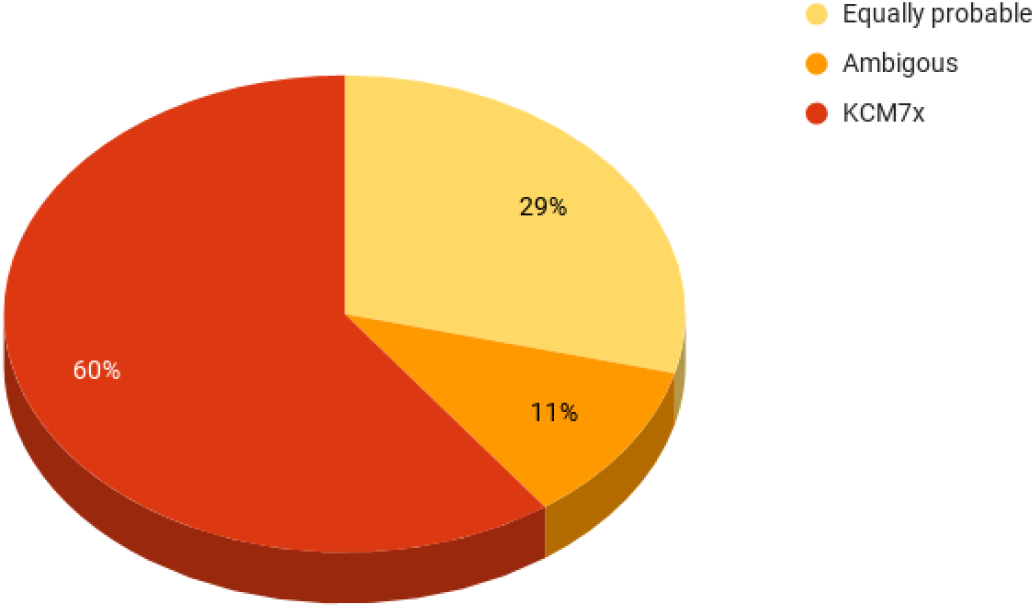
The percentage of the data sets in favor of *KCM_7x_* and *KCM_7xM0_* models for 70 empirical data sets based on Δ*AICc* measure.

**Figure 10.**
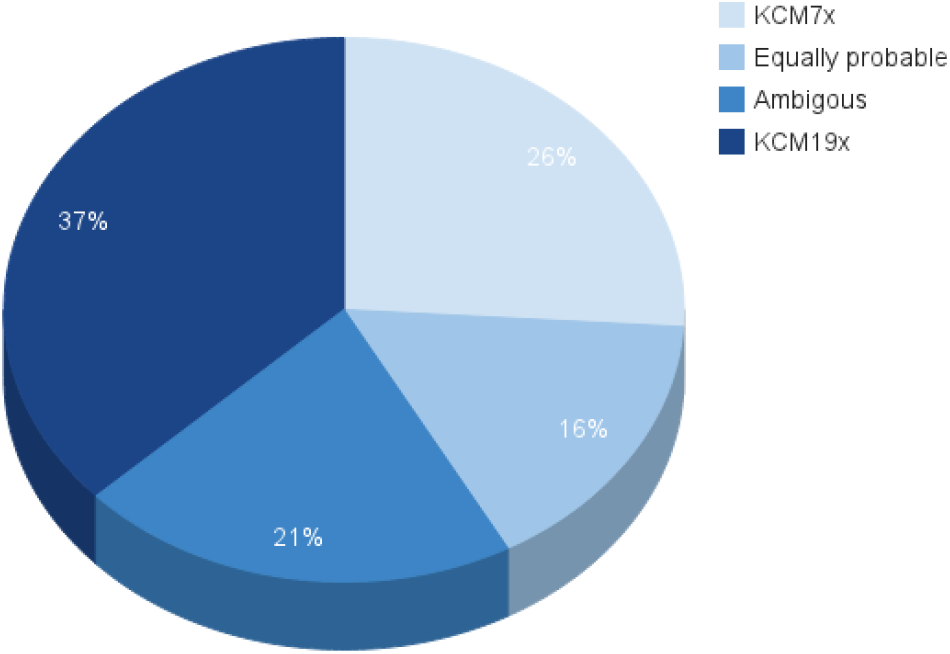
The percentage of the data sets in favor of *KCM_7x_* and *KCM_19x_* models for 70 empirical data sets based on Δ*AICc* measure.

Our results show that GTR-based models always outperform HKY-based models when double and triple substitutions are allowed. Furthermore, our observations show that the performance of the HKY-based models of the framework are decreased when they allow double and triple substitutions. In particular, *KCM_2x_* and *KCM_4x_*, perform worst than the member models that ignore double and triple substitutions, *KCM_2xM0_* and *KCM_4xM0_*. Even, when double and triple substitutions are neglected GTR-based models outperform HKY based ones (table 2). This observation is consistent with the results of [46], where the superiority of the GTR-based model over the HKY-based model was claimed while ignoring double and triple substitution on the large data sets. Trivially, the better fit of the GTR-based models over the HKY-based models is even more evident based on our results when a model allows double and triple substitution.

### Parameters

One of the advantages of mechanistic models is that their parameters are defined based on biological processes [18] and allow a direct test of the relevance of these parameters through AIC or likelihood-based measures.

Based on our results, the models of the *KCM-based model family* framework can estimate the transition-transversion bias with proper accuracy (table 4, figures 11, 3, 12). In particular, the entities of the 4 × 4 matrices used as the bases of the models of the framework estimate the mutational bias at the nucleotide level. This is true even for the models that allow double and triple substitutions, i.e. *KCM_19x_* and *KCM_7x_*. The other approach, i.e. SDT model [16], that allows double and triple substitutions in a mechanistic codon model, estimates the codon rate matrix by transition-transversion rate and selection pressure parameters. The model over-estimates transition-transversion bias and it has been mentioned that the reason is unknown to the authors. The mechanistic-empirical models that allow double and triple substitutions have the effect of transition-transversion rate in exchangability rates and the estimation of transition-transversion bias is not straight forward for these models [21].

**Table 4.** The p-value of the Wilcoxon rank sum test comparing the transition-transversion ratio, *R*, and the selection pressure, *ω*, estimated by the *KCM-based model family* models, for 70 empirical data sets

| Model | p-value for $R$ | p-value for $\omega$ |
| --- | --- | --- |
| $KCM_{4xM0}$ | 0.3559 | 0.9784 |
| $KCM_{7xM0}$ | 0.5526 | 0.8366 |
| $KCM_{7x}$ | 0.6362 | 0.2721 |
| $KCM_{19xM0}$ | 0.8268 | 0.2150 |
| $KCM_{19x}$ | 0.3826 | 0.1872 |

**Figure 11.**
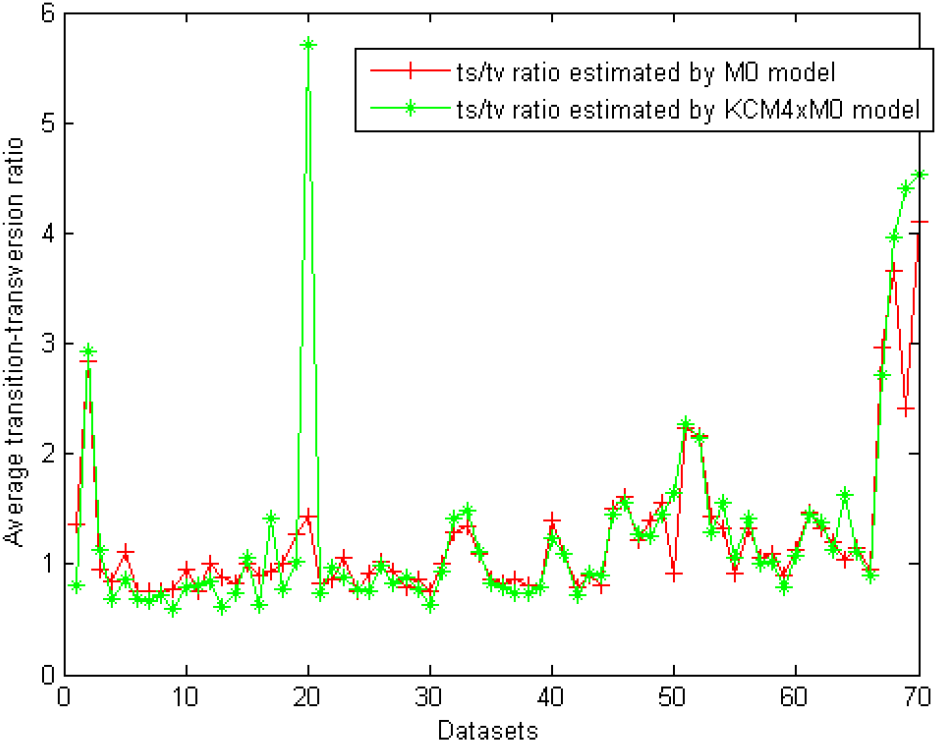
Estimation of the average rate of transition and transversion of *KCM_4xM0_* for the 70 empirical data sets selected from the Selectome database. The data sets are sorted based on the *ω* value estimated by *KCM_2xM0_* model.

**Figure 12.**
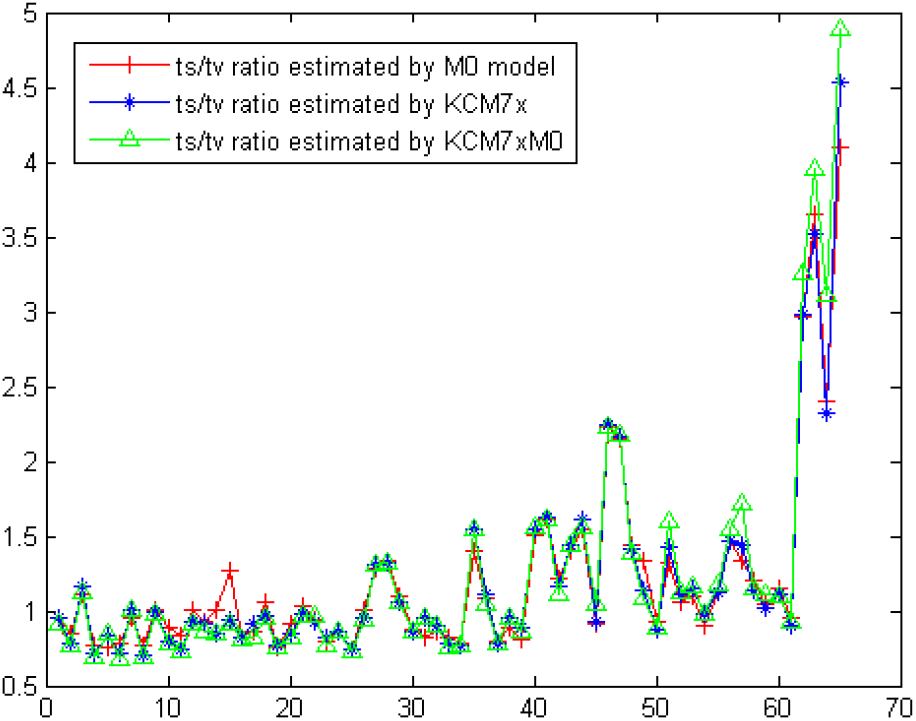
Estimation of the average rate of transition and transversion of *KCM_7x_* and *KCM_7xM0_* models for the empirical data sets selected from the Selectome database. The data sets are sorted based on the *ω* value estimated by *KCM_2xM0_* model.

**Figure 13.**
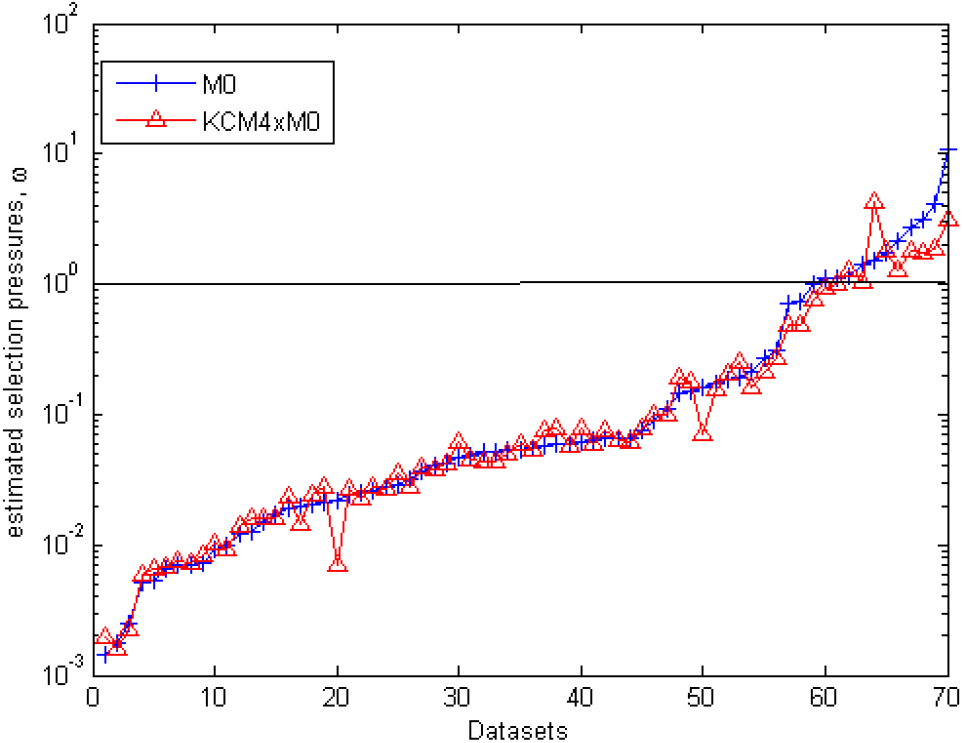
Selection pressure, *ω*, estimated by *KCM_4xM0_* and *KCM_2xM0_* models for the empirical data sets selected from the Selectome database. The data sets are sorted based on the *ω* value estimated by *KCM_2xM0_* model (y-axes is in logarithmic scale).

Regarding the selection pressure estimation, our results acknowledge the previous results that if a data set has adequate amount of double and triple substitutions, then ignoring them results to the biased estimation of the *ω* parameter [16]. Because the model that does not allow double and triple substitution break down one double or triple nonsynonymous substitution to multiple single nonsynonymous and synonymous substitutions. Accordingly, *ω* should be artificially biased to describe the double and triple substitutions in the data set [16]. When we allow double and triple substitutions, i.e. *KCM_7x_* and *KCM_19x_* models, the estimated selection pressure is lower than the one estimated by *KCM_2xM0_* in most of the data sets (figures 14, 4). Comparing to the SDT model, the only mechanistic model that allows double and triple substitutions, it estimates the selection pressure with two parameters *ω*_1_ and *ω*2. The latter is irrelevant to our model, because it shows the effect of selection pressure on two neighboring codons. The former, i.e. *ω*_1_, is comparable to the *ω* parameter estimated by the *KCM_7x_* and *KCM_19x_* models. As it is reported by [16], the *ω* estimated by SDT model under-estimate the *ω* estimated by *KCM_2xM0_* model on the empirical data set. The same pattern can be seen for *ω* estimated by *KCM_7x_* and *KCM_19x_* compare to *KCM_2xM0_* on most of the data sets.

**Figure 14.**
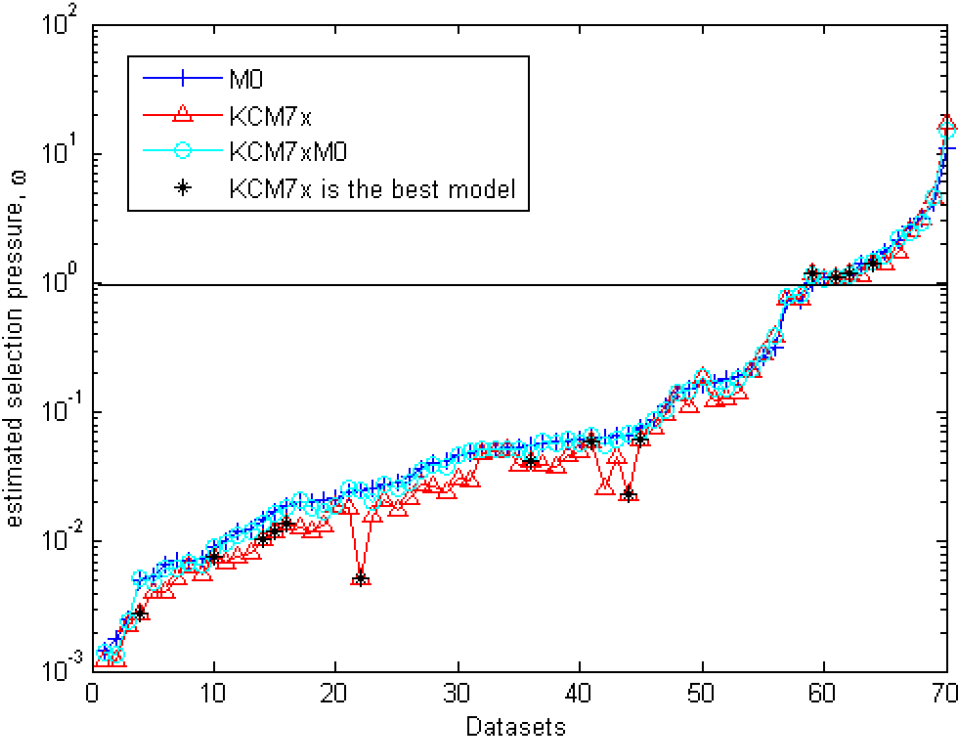
Selection pressure, *ω*, estimated by *KCM_7xM0_*, *KCM_7x_* and *KCM_2xM0_* models for empirical data sets selected from the Selectome database. The data sets are sorted based on the *ω* value estimated by *KCM_2xM0_* model (y-axes is in logarithmic scale).

Moreover, [7] has developed a model with three 4 × 4 GTR matrices for each codon position and investigated the effect of the codon frequency models on the estimation of selection pressure. They show that using *F*1 × 4, *F*3 × 4 and *F*61 frequency models [40] causes some biases on the estimation of selection pressure. Accordingly, they propose a frequency model based on conditional nucleotide frequency. Since we use *F*3 × 4 in all of the models of the *KCM-based model family* framework, the biases in the selection pressure is not because of the frequency model choice. However, we have shown that the frequency model is effective on the performance of the codon models [40]. Therefore, applying the frequency model proposed by [7] should improve the performance of the models of the *KCM-based model family* framework.

## Conclusion

As a conclusion, we developed a framework of codon based models, based on holding and relaxing the three main simplified assumptions underlying most of the current mechanistic codon models. We applied these models to empirical data sets and we compared the models of the *KCM-based model family* framework in order to choose the best model for the data set in hand. Based on the results, we showed that at least 40% of data sets need a complex model that relaxes the three simplified assumptions.

Looking at the results obtained by simulations and the results from empirical data sets, following two statements are concluded:

- Generally, models that allow double and triple substitutions perform better on empirical and simulated data sets. The *KCM_7xM0_* and *KCM_19xM0_* models never, with high evidence ratio, fit better than their alternative models that allow double and triple substitutions on simulation and empirical data sets, even when the data sets are simulated without double and triple codon substitutions. This is maybe due to the fact that the number of parameters are the same, and when a model forces double and triple substitutions to be zero, the optimization process get confused and stuck in a local minimum. However, when a model that allows double and triple substitutions fit significantly better to the data set compare to the alternative model, it shows that the amount of double and triple is not neglectable.
- The next conclusion is that the models that allow mutation heterogeneity between nucleotides within codons, i.e. *KCM_19xM0_* and *KCM_19x_* fit better than their alternative models, *KCM_7xM0_* and *KCM_7x_*, on about 40% of empirical data sets. On the other hand, on simulation results, *KCM_19xM0_* and *KCM_19x_* both fit better compare to their alternative models on 40% of data sets when the noise level is around 15%. Under the assumption that mutation heterogeneity in the nature is coming from a normal distribution, one can conclude that at least in 40% of empirical data sets the rate variation across nucleotides within a codon is equal or more than 15%.

We show that M0 model is a subset of *KCM-based model family*, and we conjecture that we can interpret original *ECM* codon substitution rate matrix as an average of substitution rate matrices of many the best fitted model of the *KCM-based model family* framework over empirical data sets, i.e. while the best fitted model is able to calculate the substitution rate matrix of individual data sets, ECM model is an average over many.

## Acknowledgments

The authors thank Maria Anisimova and Ziheng Yang for helpful discussions on this study. This work was supported by the Swiss National Science Foundation grants (3100A0-116412 and 3100A0-182312) to N.S. The Vital-IT facilities of the Swiss Institute of Bioinformatics were used for all computational aspects of this study.

## Supporting Information

### S1_ Simulation1

#### From M0 to *KCM*4xM0, *KCM*7xM0 and *KCM*19xM0

Focusing on the first class of simulations performed on the lower side (Figure 6), we start from M0 model. Recalling that the formula of M0 model in *KCM-based model family* framework is as follow: 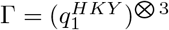, we apply two different types of noises to move towards the three other models in the same space.

- Type 1 noise (GTR noise), is applied on the entities of 4 × 4 building block of the reference model. We move from M0 model towards *KCM*7xM0 model by gradually changing the structure of the 4 × 4 building block of M0 model from HKY to GTR. For this purpose, the noises 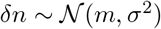, where *m* is the entity of the HKY matrix and *σ* is chosen based on the level noise, from the following set [0.05, 0.1, 0.15, 0.2], are applied to the entities of HKY matrix. Therefore the formula of *KCM*2xM0 moves towards *KCM*7xM0, i.e. 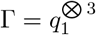 with different percentage of noises (5%, 10%, 15% *and* 20%) as shown in figure 15.
- Type 2 noise (mutation variation noise), is applied in a way to differentiate the unique 4 × 4 building blocks to the three different 4 × 4 building blocks. We move from M0 model towards *KCM*4xM0 model by changing the number of 4 × 4 building block of M0 model from one to three gradually. This is done by multiplying noises only to the transition-transversion parameter of the HKY model *κ*. The noises are drawn from 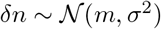 where *m* = *κ* and *σ* is from the following set [0.05, 0.1, 0.15, 0.2]. As a result of the mentioned multiplication we obtain three different 4 × 4 HKY building blocks. Kronecker product of them and the following post processes mentioned in Method section results in the codon substitution matrix of *KCM*4xM0 with the following formula: 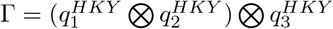.

**Figure 15.**
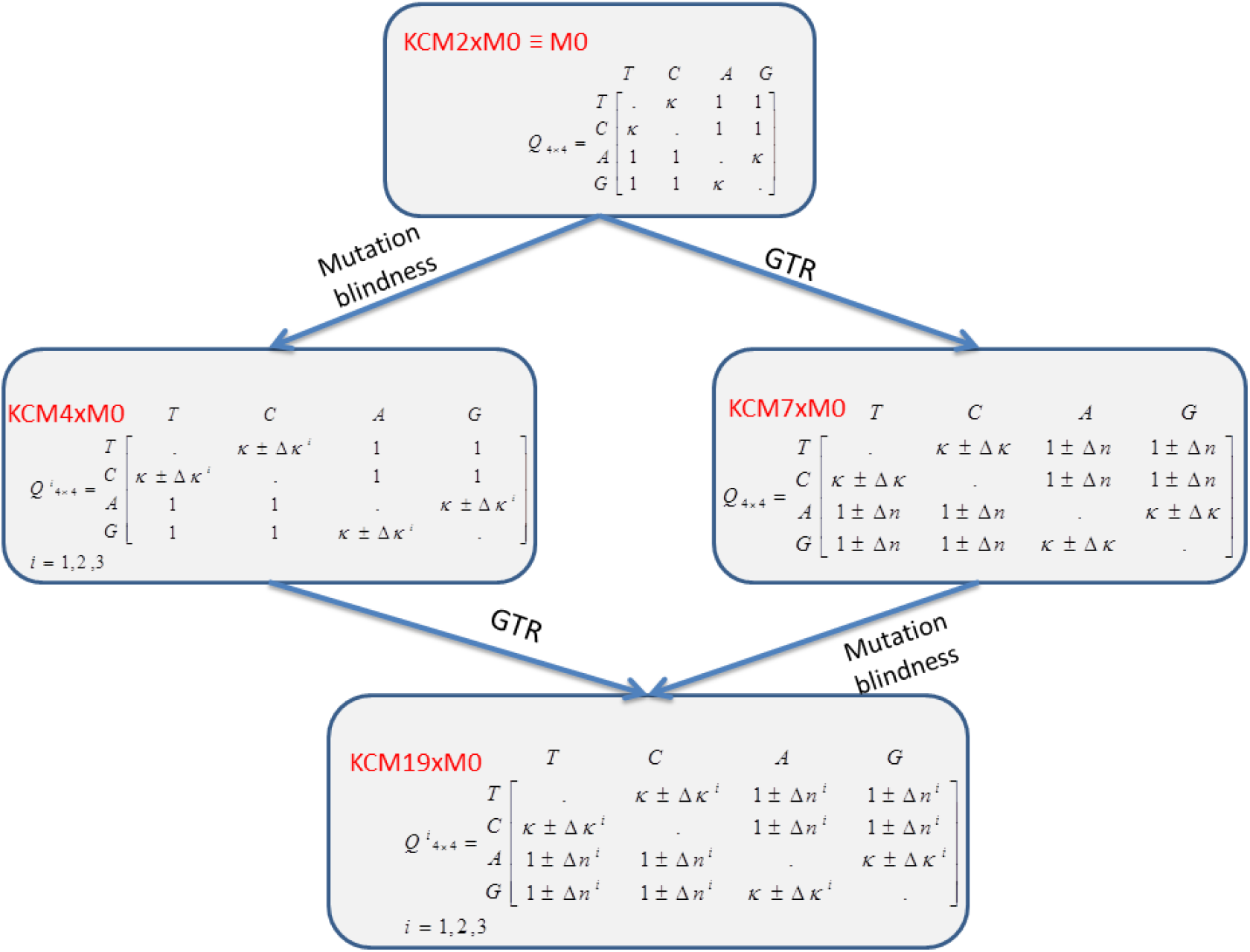
An illustration of the details of the first set of simulations.

**Figure 16.**
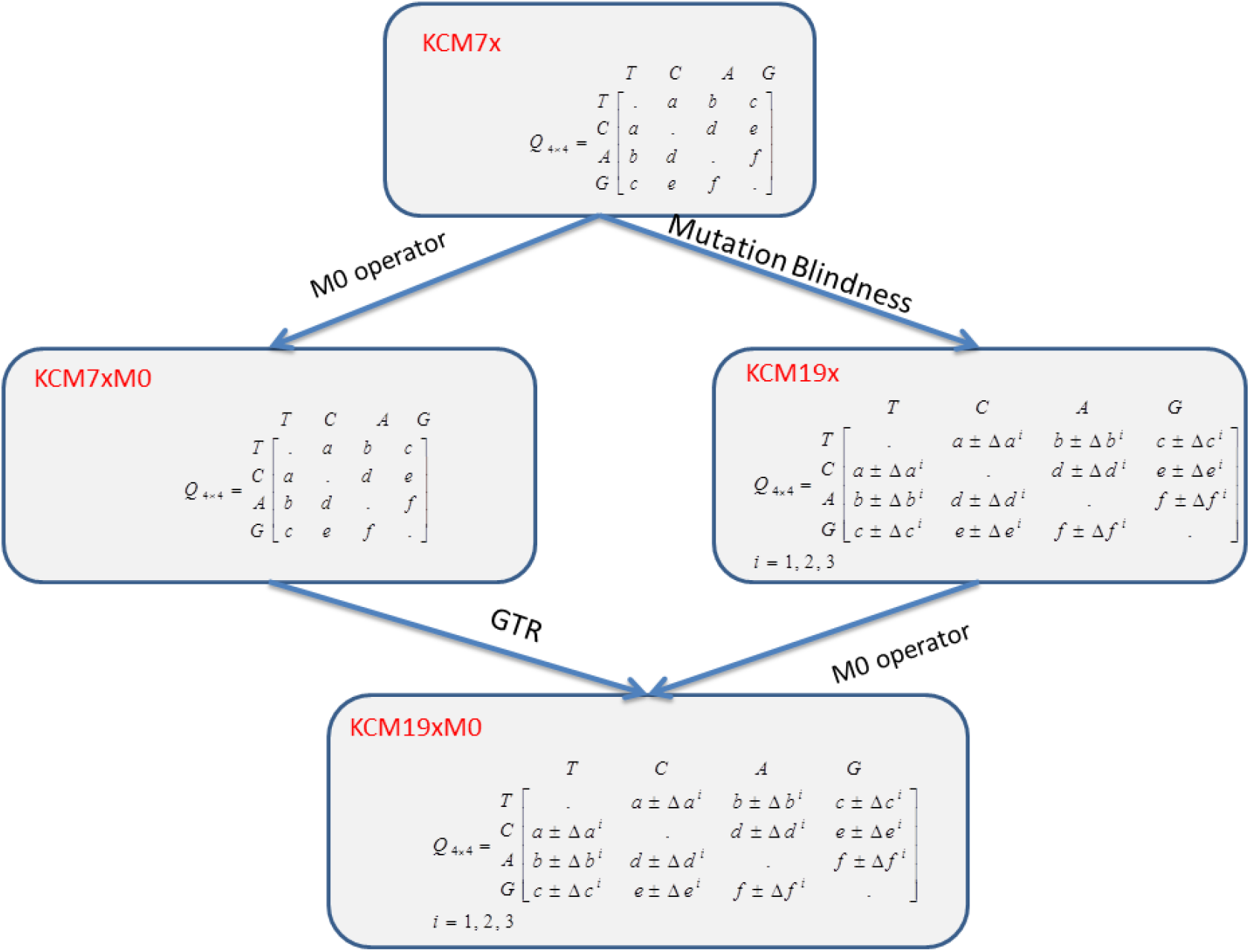
An illustrative explanation of the details of the second set of simulations.

Finally, one can move from M0 model towards *KCM*19xM0 model by applying both types of noises simultaneously, and changing the formula of M0 model to 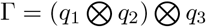.

### S2 Simulation2

#### From *KCM*7x to *KCM*7xM0, *KCM*19xM0 and *KCM*19x

The second class of simulations performed on the right side of the cubic illustration (Figure 6), we start from *KCM*7x model. In order to get to the *KCM*19x, type 2 noise, the mutation variation noise is applied to *KCM*7x with the same level of noises explained above. Moving to *KCM*7xM0 and *KCM*19xM0 is performed by introducing the type 3 noise (M0 noise) as follow:

- Type 3 noise (M0 noise), is applied directly to the codon transition rate matrix of *KCM*7x in a way to decrease the rate of double and triple substitution gradually from 25% to 6%, 4%, 3%, 1.5% and 0.05% in average. Table 5 shows the distribution of noises and the corresponding percentage of double and triple rates after applying each level of noises.

**Table 5.**
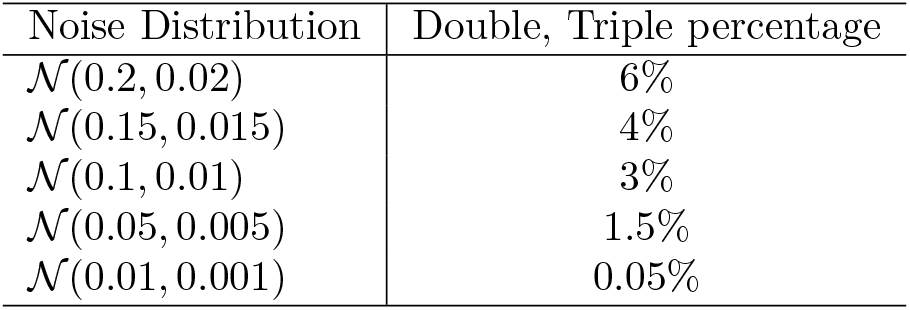
Type 3 noise, M0 noise distributions and the resulted percentage of double and triple substitutions.

**Table 6.**
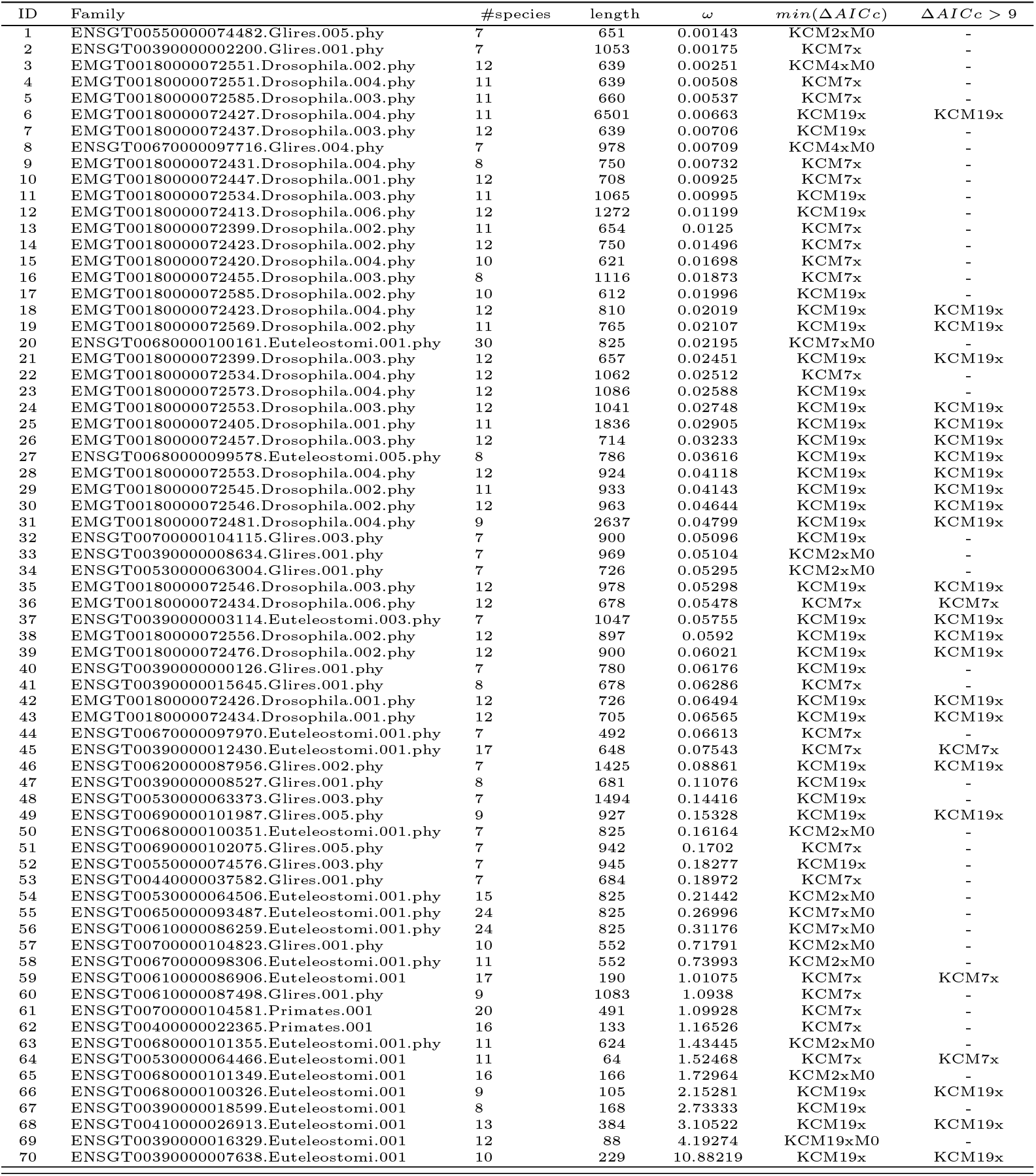
List of the 70 empirical data sets selected from the Selectome database. The value of *ω* was estimated with the *M0* model.

By applying M0 noise, *KCM*7x model moves toward *KCM*7xM0. By accumulating mutation variation noise and M0 noise simultaneously, we move toward *KCM*19xM0 model.

### Data

## Literature cited

1. Benzer S (1961) On the topography of the genetic fine structure. Genetics 47: 403–415.

2. Yang Z (1996) Among-site rate variation and its impact on phylogenetic analyses. Trends Ecol Evol 11: 367–372.

3. Hwang D, Green P (2004) Bayesian markov chain monte carlo sequence analysis reveals varying neutral substitution patterns in mammalian evolution. Proc Natl Acad Sci 101: 1399414001.

4. Gaut B, Yang L, Takuno S, Eguiarte L (2011) The patterns and causes of variation in plant nucleotide substitution rates. Annu Rev Ecol Evol Syst 42: 245266.

5. Davydov II, Salamin N, Robinson-Rechavi M (2019) Large-Scale Comparative Analysis of Codon Models Accounting for Protein and Nucleotide Selection. Molecular Biology and Evolution 36: 1316–1332.

6. Wisotsky SR, Kosakovsky Pond SL, Shank SD, Muse SV (2020) Synonymous Site-to-Site Substitution Rate Variation Dramatically Inflates False Positive Rates of Selection Analyses: Ignore at Your Own Peril. Molecular Biology and Evolution 37: 2430–2439.

7. Yap V, Lindsay H, Easteal S, Huttley G (2010) Estimates of the effect of natural selection on protein-coding content. Mol Biol Evol 27: 726–734.

8. Tian D, Wang Q, Zhang P, Araki H, Yang S, et al. (2008) Single-nucleotide mutation rate increases close to insertions/deletions in eukaryotes. Nature 455: 1058.

9. Ossowski S, Schneeberger K, Lucas-Lledo J, Warthmann N, Clark R, et al. (2010.) The rate and molecular spectrum of spontaneous mutations in arabidopsis thaliana. Science 327: 9294.

10. Goldman N, Yang ZH (1994) Codon-based model of nucleotide substitution for protein-coding dna-sequences. Mol Biol Evol 11: 725–736.

11. Nielsen R, Yang ZH (1998) Likelihood models for detecting positively selected amino acid sites and applications to the hiv-1 envelope gene. Genetics 148: 929–936.

12. Kosakovsky Pond SL, Muse SV (2005) Site-to-site variation of synonymous substitution rates. Mol Biol Evol 22: 2375–2385.

13. Seoighe C, Ketwaroo F, Pillay V, Scheffler K, Wood N, et al. (2007) A model of directional selection applied to the evolution of drug resistance in hiv-1. Mol Biol Evol 24: 1025–1031.

14. Wong WSW, Sainudiin R, Nielsen R (2006) Identification of physicochemical selective pressure on protein encoding nucleotide sequences. BMC Bioinf 7.

15. Murrell B, Wertheim JO, Moola S, Weighill T, Scheffler K, et al. (2012) Detecting individual sites subject to episodic diversifying selection. PLoS Genet 8.

16. Whelan S, Goldman N (2004) Estimating the frequency of events that cause multiple-nucleotide changes. Genetics 167: 2027–2043.

17. Dunn KA, Kenney T, Gu H, Bielawski JP (2019) Improved inference of site-specific positive selection under a generalized parametric codon model when there are multinucleotide mutations and multiple nonsynonymous rates. BMC Evolutionary Biology 19: 22.

18. Yang Z (2006) Computational Molecular Evolution. Oxford Series in Ecology and Evolution. oxford university press.

19. Cannarozzi G, Schneider A (2012) Codon evolution: mechanisms and models. Oxford: Oxford University Press.

20. Doron-Faigenboim A, Pupko T (2007) A combined empirical and mechanistic codon model. Mol Biol Evol 24: 388–397.

21. Kosiol C, Holmes I, Goldman N (2007) An empirical codon model for protein sequence evolution (vol 24, pg 1464, 2007). Mol Biol Evol 24: 2151–2151.

22. Dayhoff MO, Schwartz RM, Orcutt BC (1978) Atlas of protein sequence and structure, volume 5. National Biomedical Research Foundation, 345–351 pp.

23. Whelan S, Goldman N (2001) A general empirical model of protein evolution derived from multiple protein families using a maximum-likelihood approach. Mol Biol Evol 18: 691–699.

24. Averof M, Rokas A, Wolfe KH, Sharp PM (2000) Evidence for a high frequency of simultaneous double-nucleotide substitutions. Science 287: 1283–1286.

25. Besenbacher S, Sulem P, Helgason A, Helgason H, Kristjansson H, et al. (2016) Multi-nucleotide de novo mutations in humans. PLOS Genetics 12: 1–15.

26. Aguileta G, Bielawski JP, Yang ZH (2004) Gene conversion and functional divergence in the beta-globin gene family. J Mol Evol 59: 177–189.

27. Drake JW (2007) Too many mutants with multiple mutations. Crit Rev Biochem Mol Biol 42: 247–258.

28. Hershberg R, Petrov DA (2008) Selection on codon bias. Annu Rev Genet 42: 287–299.

29. Chuzhanova NA, Anassis EJ, Ball EV, Krawczak M, Cooper DN (2003) Meta-analysis of indels causing human genetic disease: Mechanisms of mutagenesis and the role of local dna sequence complexity. Hum Mutat 21: 28–44.

30. Smith NGC, Webster MT, Ellegern H (2003) A low rate of simultaneous double-nucleotide mutations in primates. Mol Biol Evol 20: 47–53.

31. Yang Z (1994) Estimating the pattern of nucleotide substitution. J Mol Evol 39: 105–111.

32. Nei M, Gojobori T (1986) Simple methods for estimating the numbers of synonymous and nonsynonymous nucleotide substitutions. Mol Biol Evol 3: 418–426.

33. Jukes T, Cantor C (1969) Evolution of Protein Molecules. New York: Academic Press.

34. Yang Z, Nielsen R (2000) Estimating synonymous and nonsynonymous substitution rates under realistic evolutionary models. J Mol Evol 17: 32–43.

35. De Maio N, Holmes I, Schloetterer C, Kosiol C (2013) Estimating empirical codon hidden markov models. Mol Biol Evol 30: 725–736.

36. Yang ZH, Nielsen R (2002) Codon-substitution models for detecting molecular adaptation at individual sites along specific lineages. Mol Biol Evol 19: 908–917.

37. Nielsen R, Huelsenbeck J (2002) Detecting positively selected amino acid sites using posterior predictive p-values. Pac Symp Biocomput 7: 576588.

38. Lemey P, Minin V, Bielejec Fea (2012) A counting renaissance: combining stochastic mapping and empirical bayes to quickly detect amino acid sites under positive selection. Bioinformatics 28: 3248–3256.

39. Dutheil JY, Galtier N, Romiguier Jea (2012) Efficient selection of branch-specific models of sequence evolution. Mol Biol Evol 29: 1861–1874.

40. Zaheri M, Dib L, Salamin N (2014) A generalized mechanistic model of codon evolution. Mol Biol Evol.

41. Tavare S (1986) Some probabilistic and statistical problems in the analysis of dna sequences. Lectures Math Life Sci, Amer Math Soc: 57–86.

42. Uzzell T, Corbin KW (1971) Fitting discrete probability distributions to evolutionary events. Science 172: 1089–1096.

43. Yang Z (1993) Maximum-likelihood estimation of phylogeny from dna sequences when substitution rates differ over sites. Mol Biol Evol 10: 1396–1401.

44. Yang Z (1994) Maximum likelihood phylogenetic estimation from dna sequences with variable rates over sites: Approximate methods. J Mol Evol 39: 306–314.

45. Zoller S, Schneider A (2010) Empirical analysis of the most relevant parameters of codon substitution models. J Mol Biol 70: 605–612.

46. Delport W, Scheffler K, Seoighe C (2009) Models of coding sequence evolution. Brief Bioinform 10: 97–109.

